# Evolution of thermal performance curves: a meta-analysis of selection experiments

**DOI:** 10.1101/2022.05.09.491229

**Authors:** Sarthak P. Malusare, Giacomo Zilio, Emanuel A. Fronhofer

## Abstract

Temperatures are increasing due to global changes, putting biodiversity at risk. Organisms are faced with a limited set of options to cope with this situation: adapt, disperse or die. We here focus on the first possibility, more specifically, on evolutionary adaptations to temperature. Ectotherms are usually characterized by a hump-shaped relationship between fitness and temperature, a non-linear reaction norm that is referred to as thermal performance curve (TPC). To understand and predict impacts of global change, we need to know whether and how such TPCs evolve.

Therefore, we performed a systematic literature search and a statistical meta-analysis focusing on experimental evolution and artificial selection studies. This focus allows us to directly quantify relative fitness responses to temperature selection by calculating fitness differences between TPCs from ancestral and derived populations after thermal selection.

Out of 7561 publications screened, we found 47 studies corresponding to our search criteria representing taxa across the tree of life, from bacteria, to plants and vertebrates. We show that, independently of species identity, the studies we found report a positive response to temperature selection. Considering entire TPC shapes, adaptation to higher temperatures traded off with fitness at lower temperatures, leading to niche shifts. Effects were generally stronger in unicellular organisms. By contrast, we do not find statistical support for the often discussed “Hotter is better” hypothesis.

While our meta-analysis provides evidence for adaptive potential of TPCs across organisms, it also highlights that more experimental work is needed, especially for under-represented taxa, such as plants and non-model systems.

## Introduction

Human-induced global increases in temperature have been responsible for, and projected to cause, large-scale global extinctions of species (Sinervo *et al*., 2010), range shifts (Parmesan, 2006; Root *et al*., 2003), and changes in the composition and dynamics of biological communities (Walther *et al*., 2002). This situation is likely to get worse in the near future, since, according to the 2021 IPCC report, global temperatures may increase by 4.4 °C until the end of the century (Masson-Delmotte *et al*., 2021). In order to understand the impacts of such temperature increases on ecological systems and generate appropriate model predictions, both ecological and evolutionary consequences of increased temperatures need to be comprehensively understood.

At a small scale, according to metabolic theory of ecology (MTE; Brown *et al*., 2004b), temperature directly affects the metabolic rates of organisms. Within a restricted range of temperatures specific to a focal organism, metabolism is thought to scale exponentially with increasing temperature. As a consequence, fitness-relevant components of an organism’s life-history, especially rates like fecundity, but also developmental time, to name but two examples, may show similar temperature dependence (Gillooly *et al*., 2001, 2002). These temperature effects may then cascade across levels of complexity, from individuals to populations and communities, and affect population dynamics, trophic interactions and many other ecological patterns (e.g., Brown *et al*., 2004a; Uszko *et al*., 2017).

Such changes in individual performance or fitness as a function of temperature are often referred to as thermal performance curves (TPC) or thermal niches. In general, the niche of an organism, defined as a set of components an organism requires from its environment along with the impact the organism exerts on its environment (Chase & Leibold, 2003), can be used to quantify the fitness response of an organism for a particular environmental requirement. Accordingly, Gvoždík (2018) defines the TPC as the range of body temperatures of ectothermic organisms in which there is positive population growth. The typical shape of the TPC is unimodal, with a gradual increase in fitness as the temperature increases until an optimum is reached beyond which fitness decreases rapidly and reaches the critical thermal limit beyond which growth is not possible (Huey & Kingsolver, 1993). This negatively skewed response can be explained by the underlying enzyme thermodynamics (Eyring, 1935; Schoolfield *et al*., 1981; Ratkowsky *et al*., 2005; DeLong *et al*., 2017).

While TPCs define how organisms react plastically to temperature, one can imagine that changes in temperature may impact TPCs evolutionarily if TPCs are heritable and enough variation is present for selection to act upon. Indeed, heritability and variation seem to be present as suggested by past research on the evolution of thermal physiology (for an overview see e.g. Angilletta *et al*., 2010). Based on this existing body of work, evolution in response to increased temperature has been predicted to lead to a phenomenon termed “Hotter is Better” (Angilletta *et al*., 2010) which posits that the maximal thermal performance of a genotype increases with temperature (Fig. 1A). Alternatively, a “Hotter is Wider” pattern has been described (Knies *et al*., 2009) which implies that genotypes with higher thermal optima may have wider TPCs (Fig. 1D). In general, the evolution of TPCs may exhibit constraints and trade-offs: as reviewed by Huey & Kingsolver (1993), thermal optima and thermal maxima seem linked but not thermal minima, for example. Trade-offs may constrain the relationship between height (maximal fitness) and width (e.g., generalist-specialist trade-off; Fig. 1C), but Sexton *et al*. (2017) argue against the universality of trade-offs and suggest that adaptation can affect TPC width significantly, for example. Most recently, Buckley & Kingsolver (2021) review the evolution of TPCs across a variety of taxa focusing on the importance of realistic environmental variation. The authors highlight that despite clear suggestions of evolutionary potential from experimental studies the evolution of thermal sensitivity seems often constrained which calls for further investigation.

**Figure 1:**
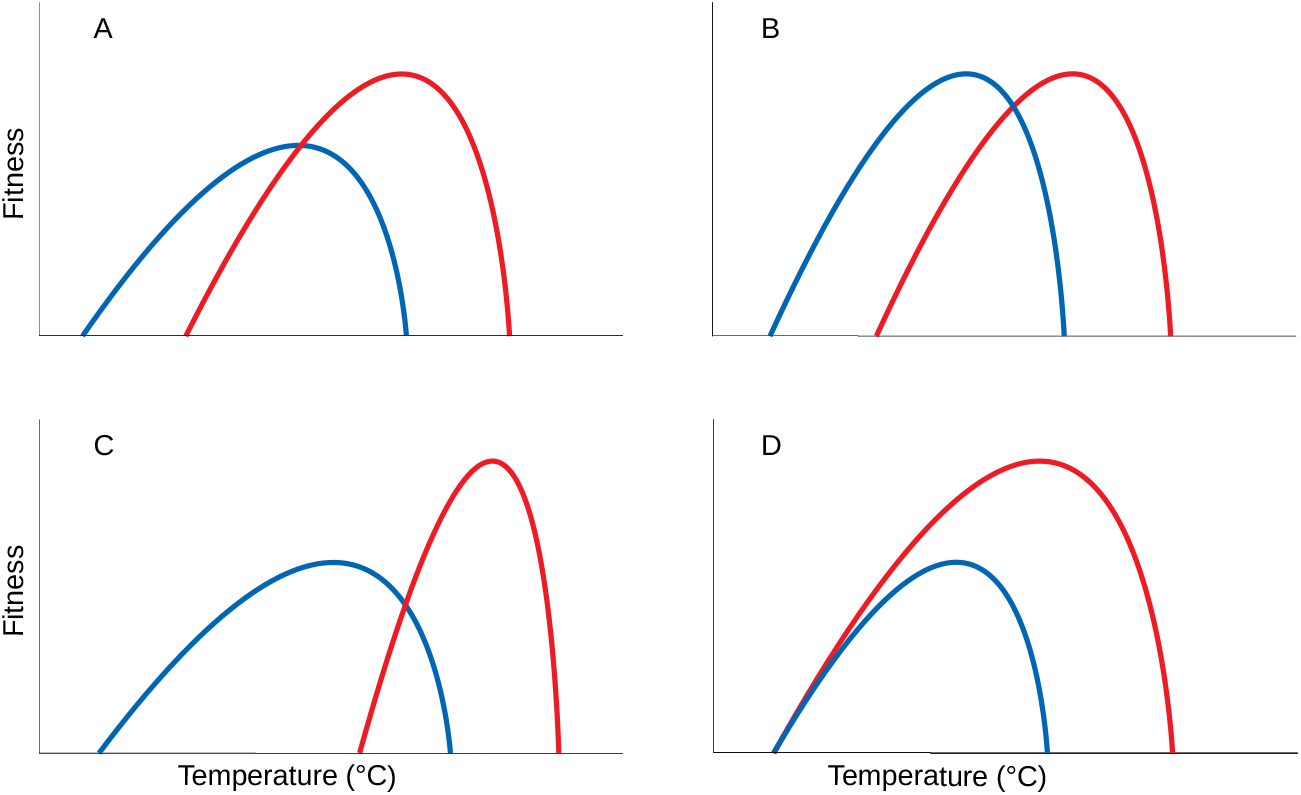
Simplified alternative hypotheses for evolutionary changes in TPCs when faced with temperature change (here only increase in temperature for simplicity; blue: ancestors; red: derived populations adapted to increased temperatures). (A) The “Hotter is Better” hypothesis (Angilletta *et al*., 2010) predicts that the maximal thermal performance of a genotype increases with temperature. (B) Alternatively, TPCs could simply shift without a change in shape. (C) Increased maximal fitness from the “Hotter is better” scenario could trade off with TPC width (generalist-specialist trade-off). Finally, (D) shows the extreme case of “Hotter is wider” (Knies *et al*., 2009) which implies a lack of constrains or trade-offs on TPC shape. Figure adapted from Knies *et al*. (2009).

Importantly, a comprehensive understanding of TPCs and their evolution is required for predicting biogeographic responses of species to climate change (Pearman *et al*., 2008; Banta *et al*., 2012; Urban *et al*., 2016; Sinclair *et al*., 2016). Furthermore, range dynamics and biodiversity maintenance is defined by evolution of TPCs which can significantly interact with other important life-history traits, such as dispersal (Hillaert *et al*., 2015; Thompson & Fronhofer, 2019). In the context of diseases, the evolution of TPCs of disease vectors critically affects disease prevalence and transmission (Corbin *et al*., 2017; Cohen *et al*., 2020; Couper *et al*., 2021; Hector *et al*., 2021).

Despite the importance of TPCs and their evolution, past synthesis work (e.g. Huey & Kingsolver, 1993; Angilletta *et al*., 2010; Buckley & Kingsolver, 2021) has remained narrative which prevents us from having a clear and especially quantitative understanding of thermal adaptation and trade-off between high and low temperature performance or other constraints. Therefore, we here carried out a systematic review and meta-analysis of TPC evolution, focusing on ectotherms (see Levesque & Marshall, 2021, for a discussion of TPCs analogs in endotherms).

In order to directly quantify relative fitness responses to temperature selection, we require TPCs measured before and after selection which is difficult to do in the field if one does not want to rely on space-for-time substitutions (transplant and common garden experiments). One possible solution are seed banks or resting stages as used by (Thomann *et al*., 2015), but we are unaware of any such study quantifying TPCs. Therefore, we focus on experimental studies that report TPCs before and after a selection or experimental evolution experiment. We quantify the relative change in fitness between ancestors and derived populations and ask whether there is a consistent evolutionary response to thermal selection across taxa, investigate potential trade-offs and analyse whether aspects such as recombination, the presence of standing genetic variation or number of generations explain evolutionary responses.

We hypothesise that adaptive evolutionary responses to temperature selection should include fitness increases, especially at the selection temperature (Fig. 1). These increases may trade off with fitness at other temperatures, although the opposite has bees suggested (“Hotter is wider” hypothesis). Finally, we will also investigate the “Hotter is better” hypothesis as described above in order to evaluate potential evolutionary constraints or the lack thereof. While our focus on experimental studies restricts the scope of our study somewhat, it allows us to quantify evolutionary changes readily. In addition to this quantitative analysis, we separately discuss work focusing on fluctuating temperature selection, on the interaction between temperature selection and biotic interactions and on thermal evolution in viruses.

## Materials and Methods

### Systematic literature search

We carried out a systematic literature search using the Web of Science database. We defined a search algorithm that allowed us to find experimental studies that performed selection on TPCs up to and including literature published in 2021 (see Supplementary Material S1 for the search algorithm; the last update to the search was conducted in April 2022 but excluded entries from 2022). We screened the outcome of this search and selected papers that had quantified a TPC of an ancestor (population from the start of the experiment) or a control (population kept at the ancestral temperature), subsequently performed a selection experiment or experimental evolution (Kawecki *et al*., 2012) and finally quantified a TPC after evolution. This allowed us to directly compare TPCs before and after evolution.

Studies included in this meta-analysis therefore had to expose populations of a study organism to a thermal condition for more than one generation. The selected papers had to contain at least one direct estimate of reproductive output like fecundity, population growth rate, fitness or another positive fitness correlate justified by the authors (see Table S1 for an overview of fitness measures used). In addition, this fitness measure had to be recorded such that it excluded purely plastic effects (e.g., via a common garden, or similar, except for studies that selected at the control temperature, i.e., did not change the environment). Papers that only contained survival could not be used because TPC shapes for birth-(usually, concave) and death-related traits (often, convex or monotonically increasing) cannot be easily compared. For simplicity, we did not consider fluctuating temperatures, biotic interactions and viruses in our meta-analysis but report qualitative results. All criteria used for rejecting papers are listed in the Supplementary Material S2 and a flow chart representation of the selection process can be found in Fig. 2.

**Figure 2:**
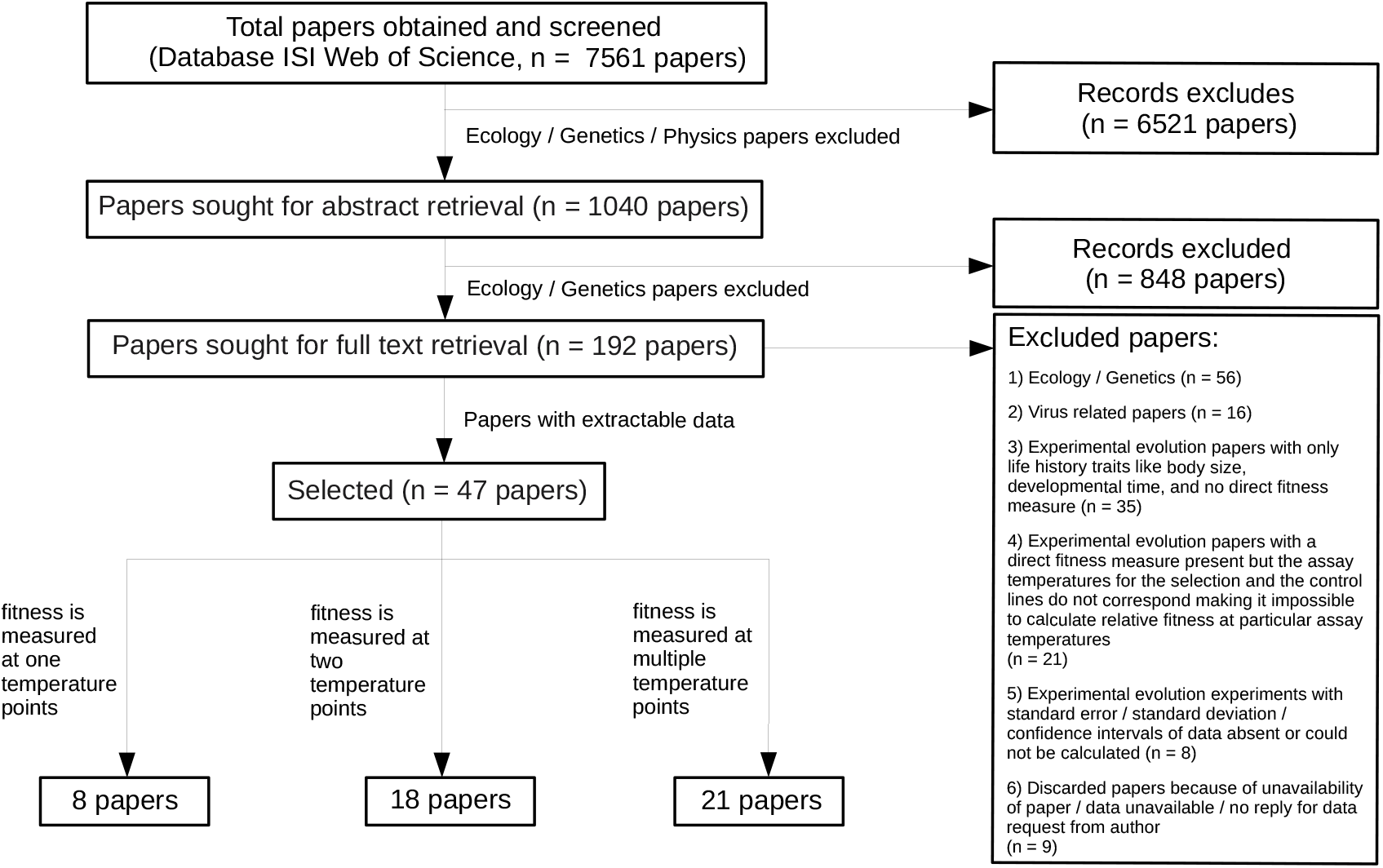
Flowchart representing the total number of papers obtained after the systematic search, along with criteria used for selecting relevant papers for the meta-analysis. The search covered all papers available in ISI Web of Science with our search criteria up to and including the year 2021 (the last update to the search was conducted in April 2022 but excluded entries from 2022).

### Data extraction

After search, screening and selection we extracted relevant data from the remaining papers, including selection temperatures, assay temperatures and fitness measures. We also recorded the number of generations an experiment lasted, the type of genetic variation used (standing genetic variation vs. de novo variation based on mutations), mode of reproduction (asexual vs. sexual; recombination may allow populations to reshuffle variation and adapt more quickly; Otto & Lenormand 2002; Moerman *et al*. 2020), whether selection was carried out at lower or higher temperatures than the ancestral/ control population and whether the reference was the ancestor or a control treatment. In addition, genus and species of the study organism was recorded, which was used to build a phylogenetic tree with the ‘taxize’ package version 0.9.99 (Chamberlain & Szocs, 2013) in R version 4.1.2 (R Core Team, 2022) and the NCBI database as a reference.

We then extracted the fitness measure as a function of temperature either directly from the papers, their supplementary information, published data associated with the paper, author supplied data or from the graphs. For extracting data from graphs we used the web-based ‘WebPlotDigitizer’ tool version 4.2 (Drevon *et al*., 2017; Rohatgi, 2020). An overview of the different methods used for different papers can be found in the Supplementary Material Table S1. This allowed us to broadly classify papers in three categories: First, studies that had selected organisms at a certain temperature and assayed fitness before and after only at this one temperature. Second, studies that include two assay temperatures yielding a classical 2-point linear reaction norm. Third, studies that had assayed ancestral/ control populations and evolved populations at more than two temperatures allowing us to study TPC shapes in more detail.

### Relative fitness calculation

In order to determine whether and how individual TPCs had evolved, we calculated fitness of the derived line relative to the ancestor or control for all assayed temperatures *T*. Relative fitness was calculated following Chevin (2011). Specifically, for taxa with discrete, non-overlapping generations we calculated relative fitness as 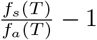 where *f*_*s*_(*T*) is fitness of the selected line at temperature *T* and *f*_*a*_(*T*) is the fitness of the ancestor or control line at the same temperature (Eq. 2.5 in Chevin (2011)). For taxa with continuous growth, we calculated relative fitness as (*f*_*s*_(*T*) − *f*_*a*_(*T*)) *G* where *G* is the generation time (Eq. 3.1 in Chevin (2011); multiplication with generation time allows us to compare studies using different time scales) and for taxa that reproduce by binary fission, we used the approximation 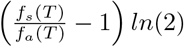 (Eq. 3.2 in Chevin (2011)). An overview can be found in the Supplementary Material Table S1.

Relative fitness was calculated using reported means over replicates. In order to take into account the associated uncertainty, we calculated the corresponding standard error (*δ*) of relative fitness assuming uncorrelated errors which, for non-overlapping generation systems gives 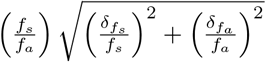. For systems with organisms reproducing by binary fission, the propagated error was calculated in analogy but multiplied with *ln*(2). For taxa with continuous growth where we did not use the binary fission approximation, the propagated relative fitness error is 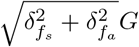. If a paper reported standard deviation instead of error we calculated the standard error by dividing the standard deviation with the square-root of the sample size. If confidence intervals were reported we calculated the standard error by dividing the confidence interval with 1.96.

### Statistical meta-analysis

The statistical meta-analysis was conducted in R version 4.1.2 using Bayesian linear mixed models that take into account measurement error (for details see Supplementary Material S3). By taking into account measurement error (calculated as described above) we effectively weighted the effect sizes (here relative fitness) by study precision (McElreath, 2020, chapter 15). The analyses were performed using the ‘rstan’ (version 2.21.3; Stan Development Team 2022) and ‘rethinking’ (version 2.21; McElreath 2020) R packages.

We excluded three of the 814 relative fitness measures extracted from a total of 47 selected papers. Two of these three excluded data points were identified as outliers with relative fitness estimates > 20 (overall mean relative fitness: 0.13). The third excluded point had an error estimate of zero and could not be included in the statistical framework used. Of the remaining 811 relative fitness measures, 60 did not report any error measurements and were also excluded.

After these data cleaning steps, we checked whether our explanatory variables (type of genetic variation used in the experiment, Var; mode of reproduction, Rep; fitness comparison, Comp; and number of generations an experiment lasted, Gen) were correlated. While the categorical explanatory variables (Var, Rep and Comp) were all only weakly corrected with the number of generations an experiment lasted (Gen; *r*(*Rep, Gen*) = 0.23, *r*(*V ar, Gen*) = 0.30, *r*(*Comp, Gen*) = 0.34; biserial correlation using the R package ‘ltm’ version 1.2; Rizopoulos 2006), the categorical variables were very strongly correlated with each other: *r*(*Rep, V ar*) = 0.98, *r*(*Rep, Comp*) = 0.80 and *r*(*V ar, Comp*) = 0.92 (tetrachoric correlation using the R package ‘psych’ version 2.2.5; Revelle 2022). As a consequence, we performed all analyses using only the number of generations an experiment lasted (Gen) and the type of genetic variation used (standing genetic variation vs. de novo variation based on mutations; Var) as explanatory variables.

We conducted four main analyses (for details see Supplementary Material S3): First, we asked whether there was an overall direct response to selection by considering fitness assays measured at the same temperature as the selection regime (230 data points). A total of 29 of the 230 TPCs assayed only this condition, whereas the remaining TPCs also tested the evolved organism at different temperatures (two-point and multipoint analysis, see below).

Second, we considered TPCs (122 data points in 61 TPCs) where organisms were assayed at two temperatures before and after selection (Two-point analysis). Here and in the following we analysed relative fitness as a function of relative assay temperature in order to be able to compare between studies using different selection temperatures. Relative assay temperature is here defined as the difference between assay and selection temperature, positive values indicating an assay temperature above the selection temperature.

Third, we analysed the TPCs (600 data points in 159 TPCs) that contain fitness at three or more assay temperatures before and after selection (Multipoint analysis). This analysis mainly allows us to study effects of thermal selection on TPC shape and potentially find a signature of the “Hotter is wider” hypothesis.

Fourth, for the TPCs that assayed more than three temperature points in the final assay (27 TPCs), we tested the “Hotter is better” hypothesis by taking the maximal fitness value of the selected and control lines to calculate relative fitness. If the “Hotter is better” hypothesis is true, relative fitness should be positive when comparing the warm-evolved TPC to the cold-evolved TPC.

In all analyses, we analysed the effects of additional explanatory variables identified from the literature that may explain variation in fitness responses. These explanatory variables were: type of genetic variation used (standing genetic variation vs. de novo variation based on mutations) and the number of generations the experiment lasted, as the other extracted variables were correlated as reported above. Note that not all studies reported the number of generations (or doubling-time periods) an experiment lasted. Therefore, we performed all analyses twice: once with the full data set, not analysing the number of generations and once with a reduced data set including the analysis of the effect of the number of generations.

We compared and ranked all models using the Watanabe-Akaike information criterion (WAIC), a generalised version of the Akaike information criterion (McElreath, 2020). We consistently used a chain length of 30,000 iterations (1,000 warmup iterations) and vaguely informative priors (for details see Supplementary Material S3).

Random effects were ‘species ID’ (see Fig. 3 and Supplementary Material S1) and ‘study ID’ in order to take species effects, as well as within study replication, into account. For the two-point and multipoint analysis we also included ‘TPC ID’ in order to avoid pooling data from independent TPC measurements.

**Figure 3:**
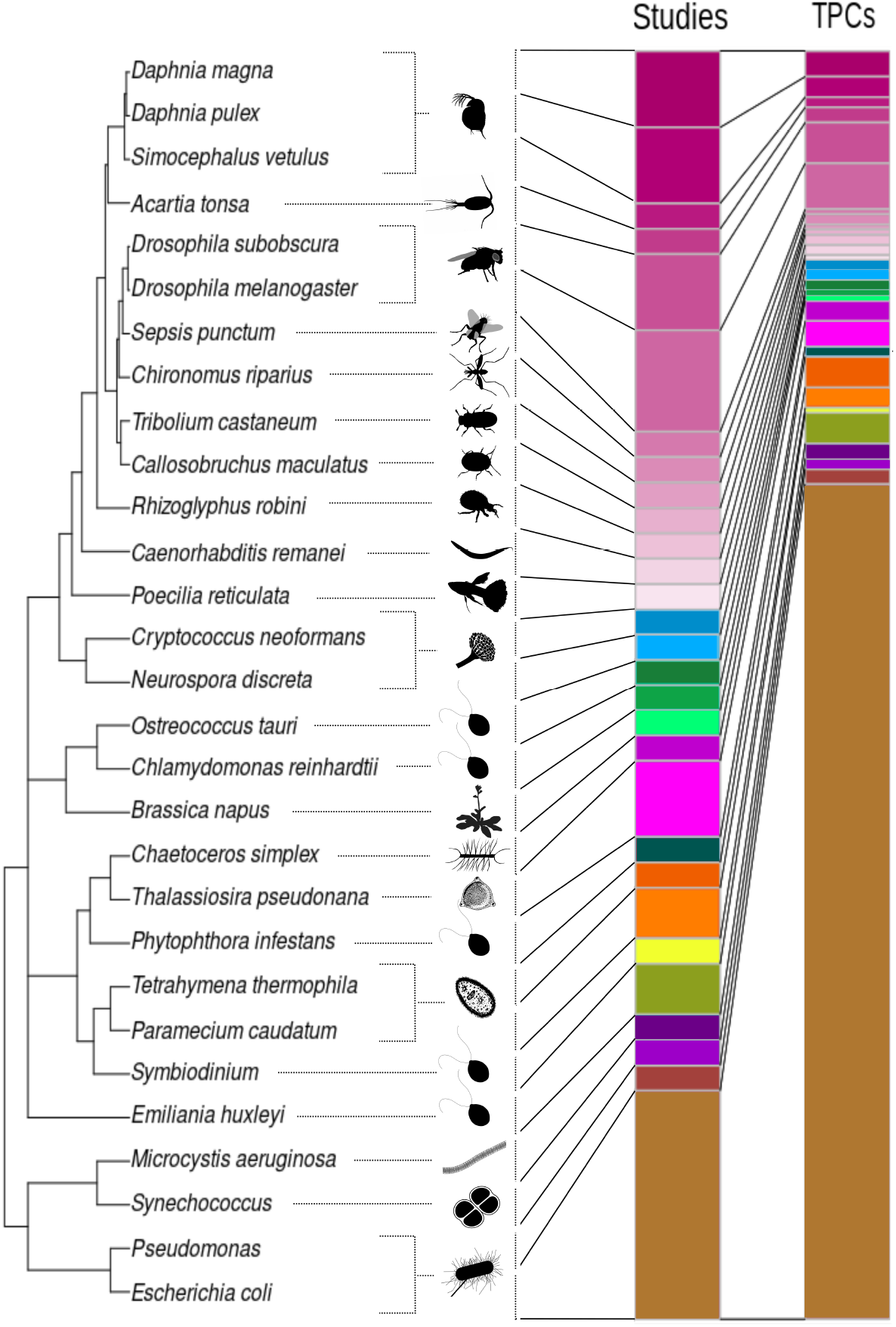
Overview of all the organisms included in the meta-analysis represented by different colours. The stacked bars represent the number of studies and the number of TPCs associated with each organism. Quantitative information can be found in Table S2.

Note that apart the ‘species ID’ random effect we did not include the phylogeny explicitly in our analyses since Cinar *et al*. (2021) show that this can be problematic when phylogenetic relationships are weak. Indeed, in our case (see Fig. 3) the phylogenetic relationships are weak with a mean correlation of our phylogenetic matrix of 0.28. We validated this conclusion by calculating the explained variance of a true phylogenetic random effect using the ‘metafor’ R package (version 3.4; Viechtbauer 2010) which, as expected, turned out to be null.

Heterogeneity was analysed using posteriors of the variances in true effects (*σ*^2^) which we can directly extract from our models for all levels, as well as *I*^2^ (Higgins & Thompson, 2002; Nakagawa & Santos, 2012). In the context of multilevel models with moderators as here, *I*^2^ can be calculated for each random effect. It captures the relative amount of variance due to true effects versus sampling error once the fixed effects are taken into account. We calculated *I*^2^ following Viechtbauer (2010) with full posteriors which easily allows us to calculate means and 95% compatibility intervals.

Finally, publication bias was assessed using funnel plots and corresponding mixed effects models (see e.g. O’Dea *et al*., 2021) which indicated no consistent bias (Figs. S1–S4). Detailed statistical methods, including information on priors can be found in the Supplementary Material S3.

## Results

From a total of 7561 publications screened, we retained 47 papers which had assayed fitness before and after thermal selection (see Fig 2). Out of these 47, eight papers had assayed their experimental lines only at the selection temperature, 18 papers examined the evolutionary response at two assay temperatures following a classical, linear reaction norm approach, and 21 papers analysed evolutionary changes in TPC shape in more detail quantifying fitness at three or more temperatures.

Although experiments with established model systems like *Escherichia coli* or *Drosophila* sp. dominated the available literature, we found papers reporting results from species that span the tree of life, from bacteria to vertebrates (Fig. 3), with a total of 29 species (see Table S2). Note that the total taxonomic breadth was even larger and also included viruses, for example. Studies on viruses, studies including biotic interactions and well as fluctuating temperatures were not included in our meta-analysis, but are discussed qualitatively below.

### Fitness response at the selection temperature

In order to evaluate whether there was an overall response to thermal selection, regardless of the taxon and other system-specific details, we analysed the fitness response of populations which were subject to a specific selection temperature at this very same temperature (data subset with 230 relative fitness estimates; 40 papers and 28 species). Estimating a global mean effect (intercept model), we found an overall positive response to selection across all 28 species included in this analysis (rel. fitness = 0.18 (0.06, 0.32); here and in the following we report medians and 95% compatibility intervals of the relevant posterior distributions).

After model selection, the best model included an effect of the type of genetic variation used in the experiments, that is, whether experiments included standing genetic variation at the beginning for selection to act upon or relied on de novo mutations (Table S8; Fig. 4A). Experiments that relied on de novo mutations yielded a stronger response to selection (rel. fitness = 0.32 (0.15, 0.52)) than experiments that started with standing genetic variation (rel. fitness = 0.05 (−0.1, 0.21)).

**Figure 4:**
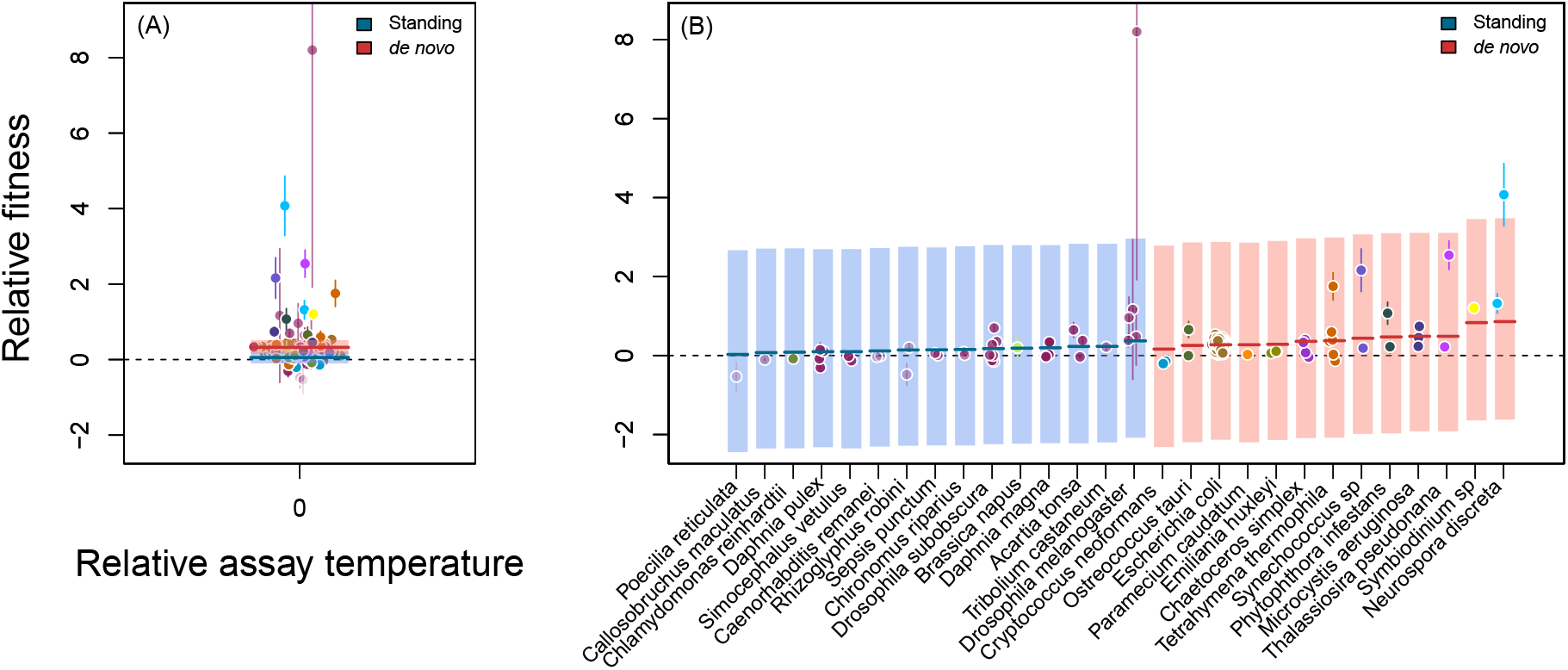
Overall response to thermal selection. (A) Relative fitness, that is, fitness of the selection lines relative to the corresponding control or ancestor assayed at the same temperature as experienced during selection (relative assay temperature of zero). Each point corresponds to one experimental populations (data subset with 230 relative fitness estimates; 40 papers and 28 species) and the error bars represent the associated standard error. Different colours correspond to different species as shown in Fig. 3. Horizontal lines visualize median posterior model predictions and shaded areas are 95% compatibility intervals. Here, the best model (Table S8) retains an effect of the type of genetic variation used in the experiments: relative fitness is higher for studies that relied on de novo mutations (red) than those that relied on selection from standing genetic variation (blue). (B) Visualisation of the ‘species ID’ random effect.

These differences were associated with study species as can be seen in Fig. 4B. Note that different studies within species did never differ in the type of genetic variation used in our data set.

Interestingly, the fitness response was independent of whether selection was for adaptation to increased or decreased temperatures since relative selection temperature was not retained during model selection (Table S8). The same was true for the number of generations an experiment lasted (Table S9).

For this overall analysis, total heterogeneity 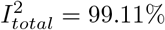 (mean; 95% compatibility interval: 98.26, 99.58) indicating that most variation was due to between species or between study heterogeneity and not to sampling error. Taking a closer look, 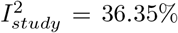 (9.97%, 94.42%) was smaller than 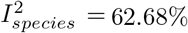 (4.32%, 89.40%). Between species heterogeneity (variance: 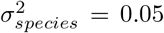 (0.003,0.14) versus 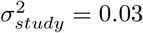 (0.01, 0.08)) can also be seen visually in Fig. 4B.

### Linear thermal reaction norms

Often, thermal selection experiments do not only assay fitness at the same temperature as selection was performed, but also add a different temperature to their assay in order to infer costs of adaptation and trade-offs, for example. Considering this subset of papers (122 data points in 61 TPCs from 18 papers and 15 species), we did not find any consistent response to selection (Fig. 5, Table S10), in contrast to the results reported in Fig. 4. None of the additional explanatory variables was retained in any of the analyses (Tables S10 and S11) and the estimated intercept overlapped with zero (rel. fitness = 0.08 (−0.04, 0.20)).

**Figure 5:**
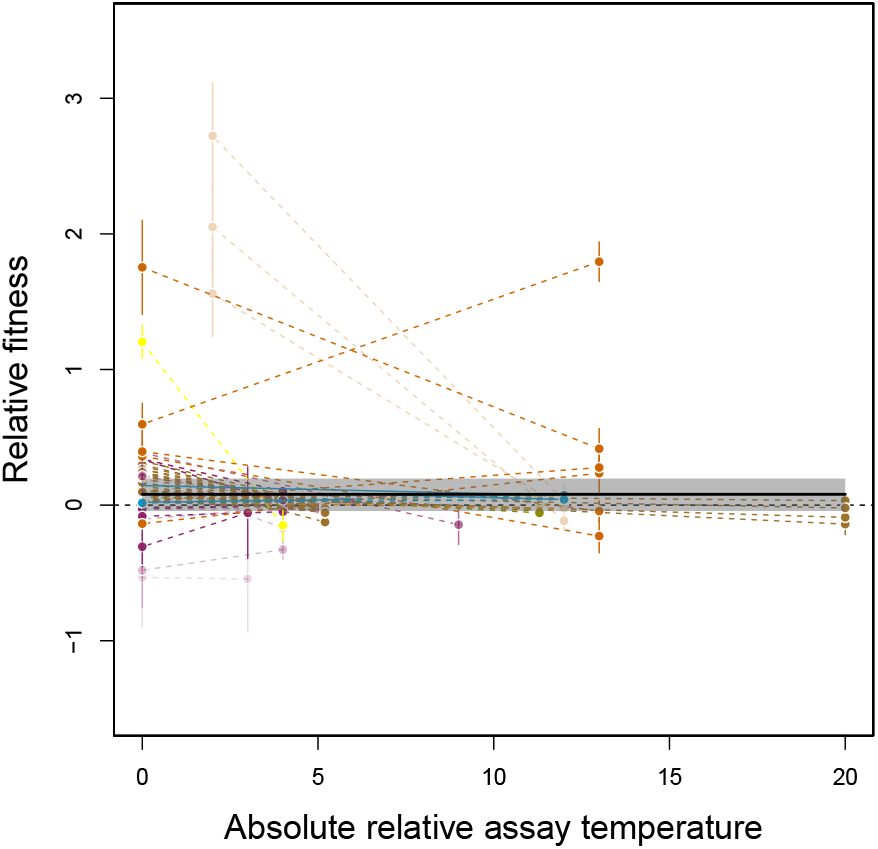
Response to selection in experiments with two assay temperatures (data subset with 122 data points in 61 TPCs from 18 papers and 15 species). The plot shows relative fitness as a function of absolute relative assay temperature, that is, the absolute difference between selection and assay temperature. Each combination of two points connected by a dashed line corresponds to one TPC (reaction norm) and the error bars represent the associated standard error. Different colours correspond to different species as shown in Fig. 3. The horizontal lines visualizes the median posterior model prediction and the shaded area is the 95% compatibility interval. Here, the best model (Table S10) is the null model and the prediction interval overlaps consistently with 0, indicating no effect of selection.

For this subset of data, heterogeneity behaved similarly to the analysis of the overall effect above. 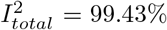 (98.08, 99.82) was very large with more heterogeneity due to species (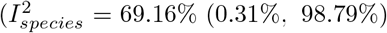; 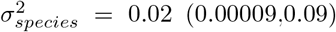 (0.00009,0.09)) than to study (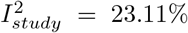 (0.05%, 97.23%); 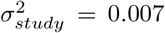 (2e-05,6e-02)) or curve (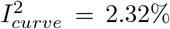 (0.005%, 29.05%); 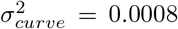 (2e-06,9e-03)).

### Evolution of the TPC shape

Studies that included more than two assay temperatures (600 data points in 159 TPCs from 21 papers and 13 species) showed a differed picture than their two-point counterparts (Fig. 6). The best fitting model retained a non-linear, cubic relationship between relative fitness and relative assay temperature (Table S12). While this specific shape should not be interpreted in any mechanistic sense, it allows us to show that while fitness is gained at and beyond the selection temperature (the 95% compatibility interval of the prediction does not overlap the zero line; see inset of Fig. 6; at a relative assay temperature of 0 and for studies that rely on de novo mutations, rel. fitness = 0.11 (0.01, 0.21)), fitness is on average lost below the selection temperature (Fig. 6) across all study species.

**Figure 6:**
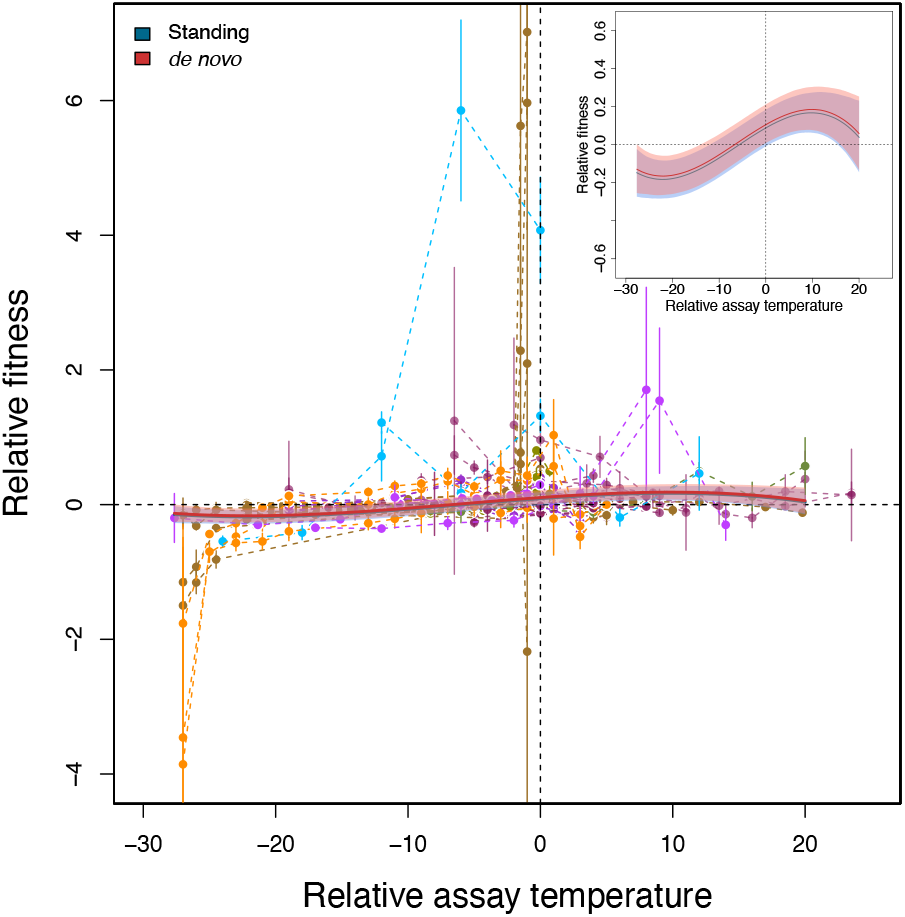
Evolution of the TPC shape (data subset of studies that included more than two assay temperatures: 600 data points in 159 TPCs from 21 papers and 13 species). The plot shows relative fitness as a function of relative assay temperature, that is, the difference between selection and assay temperature, for studies that assayed selected lines at three and more temperatures allowing us to infer changes in TPC shape. Each combination of points connected by a dashed line corresponds to one TPC and the error bars represent the associated standard error. Different colours correspond to different species as shown in Fig. 3. The solid lines visualize the median posterior model predictions and the shaded areas are the 95% compatibility interval. Here, the best model (Tables S12 and S13) is a cubic model model that includes an effect of the type of genetic variation used in the experiments (standing genetic variation vs. de novo mutations; represented in blue and red, respectively). The inset shows only the statistical model prediction for better visibility of the prediction intervals. Note that we also find an effect of relative selection temperature (‘Sign’) which we detail in Fig. S6. For simplicity, we here only show results for selection at a higher temperature compared to the control (panel C in Fig. S6). Generally, fitness tends to be gained at the selection temperature and slightly above and lost below the selection temperature and for very high temperatures.

Our analysis retained an effect of the type of genetic variation used in the experiments (standing genetic variation vs. de novo mutations; Table S12 and S13) in analogy to the overall fitness response shown in Fig. 4. Relative fitness was generally higher in experiments relying on de novo mutations (Fig. 6), although this effect was quantitatively very weak and the compatibility interval of the estimate overlapped with zero (increase in rel. fitness in experiments relying on de novo mutations: 0.01 (−0.11, 0.16)).

In addition, we found an effect of the relative selection temperature, that is, whether selection happened at higher, lower or equal temperatures compared to the control treatment (‘Sign’ in Tables S12 and S13). While the fitness response did not change qualitatively depending on ‘Sign’ (see Fig. S6; the predictions are analogous; rel. fitness difference equal-higher: 0.07 (−0.06, 0.19), rel. fitness difference equal-lower: −0.03 (−0.16, 0.11), all compatibility intervals overlap zero) studies with different relative selection temperature seem to cover different relative assay temperatures (Fig. S6).

Here, heterogeneity behaved somewhat differently in comparison to the above analyses. While 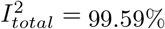 (99.11, 99.84) remained very large, most heterogeneity was due to study (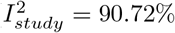 (50.87%, 99.18%); 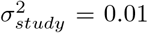 (0.006,0.035)) and not to species (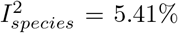 (0.01%, 47.08%); 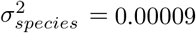 (2e-06,1e-02)) or curve (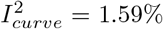 (0.004%, 10.49%); 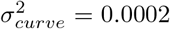 (7e-07,2e-03)). Note that while these values indicate relative changes (*I*^2^), in absolute terms (*σ*^2^) it is between species variation that is lost in this data subset.

### Is hotter better?

Finally, for the TPCs that assayed more than three temperature points, we tested the “Hotter is better” hypothesis by taking the maximal fitness value of the selected and ancestor/ control lines to calculate relative fitness (27 data points from 11 papers and 13 species). If the “Hotter is better” hypothesis is true, relative fitness should be positive when comparing the warm-evolved TPC to the cold-evolved TPC. As visible in Fig. 7, while there is a trend in the expected direction, given our dataset, we cannot confirm the “Hotter is better” hypothesis (rel. fitness = 0.08 (−0.05, 0.24)). Other explanatory variables were not retained after model selection (Tables S14 and S15).

**Figure 7:**
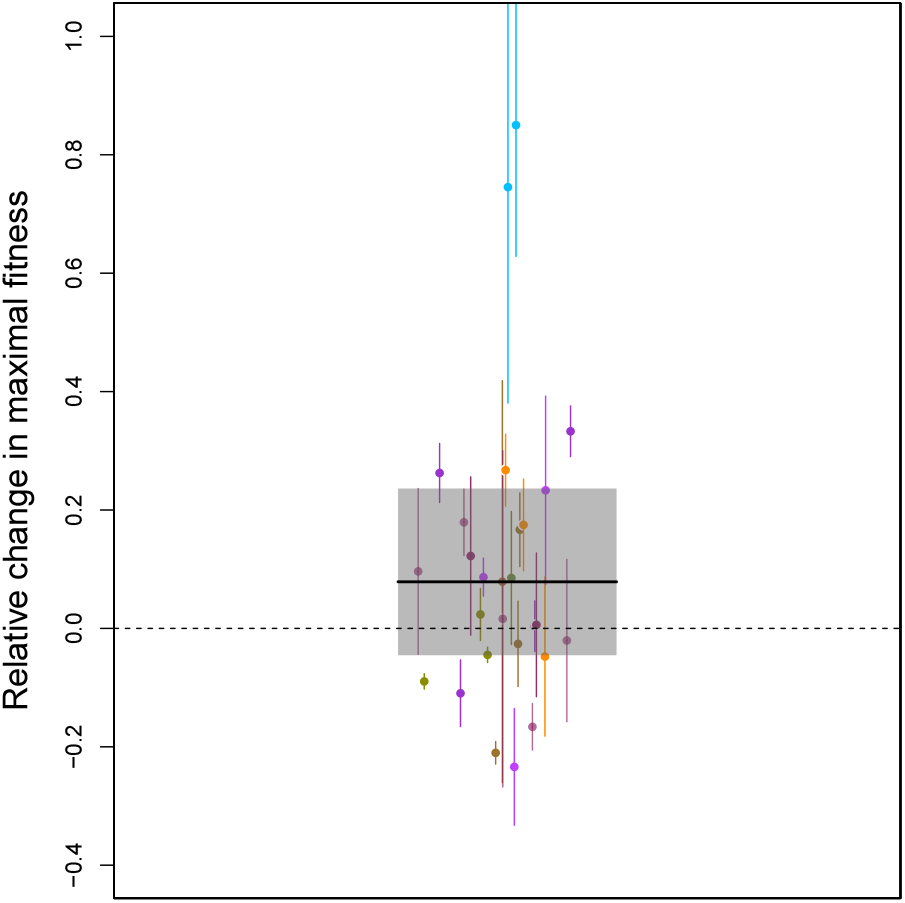
Evaluation of the “Hotter is better” hypothesis (data subset with 27 data points from 11 papers and 13 species). The plot shows relative fitness taking the maximal fitness value of the selected and ancestor/ control lines. Each points corresponds to one comparison and the error bars represent the associated standard error. Different colours correspond to different species as shown in Fig. 3. The solid line visualizes the median posterior model prediction and the shaded area is the 95% compatibility interval.

For this subset of data, heterogeneity was roughly equally distributed between random effects. 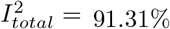 (25.87, 99.81) remained very large while heterogeneity due to species (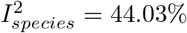 (0.07%, 94.48%); 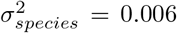 (1e-05,1e-01)) and study (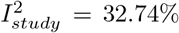 (0.07%, 94.48%); 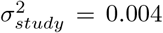 (1e-05,7e-02)) were in the same order of magnitude.

## Discussion

Our systematic literature review yielded a relatively small number of studies (47 in total) that have investigated the evolution of TPCs experimentally. While the studies covered a relatively wide range of organisms, from bacteria to plants and vertebrates (Fig. 3), a large proportion of selection experiments has been carried out on classical model organisms like *E. coli* and *Drosophila* sp. (see also Kellermann & van Heerwaarden, 2019).

Clearly, when interpreting our results one must keep in mind that we here focus on experimental evolution studies and do not include field-based comparative studies that are much more abundant and have, for instance, very recently been discussed by Buckley & Kingsolver (2021). Of course, our focus on experimental studies allows us to quantify evolutionary changes more readily since we do not rely on a space-for-time substitution, for example.

### Thermal selection leads to a consistent, positive response across taxa

Quantifying the changes in fitness due to thermal selection overall and specifically at the respective selection temperature applied in each study, we find that, across all taxa, there was a clear and positive response to selection (Fig. 4A). In addition to this global effect, we show that responses to thermal selection depended on the source of genetic variation used in the experiments, that is, whether variation was present at the beginning of the study (standing genetic variation), or whether the studies relied on de novo mutations for their evolutionary response (Fig. 4A). Somewhat counter-intuitively at first sight, studies that relied on de novo mutation showed a clearer and quantitatively stronger effect.

While we can only speculate about the underlying reasons, it is important to keep in mind that other explanatory variables correlated very strongly with the source of genetic variation. Studies that rely on de novo mutations also imply that organisms were usually asexual (correlation: *r* = 0.98). In addition, these studies tended to compare their selection outcome to an ancestor and not to a control (correlation: *r* = 0.92). The latter can imply that effects are larger because standard laboratory selection on growth, for example, due to the specific experimental setup, is not accounted for.

Independently of correlations with other explanatory variables the standing genetic variation used in the respective studies may of course not have been large enough to ensure a clear effect. However, while we do take species ID into account as a random effect, we do not include the phylogeny explicitly because our phylogenetic relationships are too weak (Cinar *et al*., 2021) due to a partially unresolved phylogeny (Fig. 3). It is therefore interesting to note that, as one can see in Fig. 4B, the reliance on de novo mutation is associated with specific species in our data set. Along the same lines, a large part of heterogeneity in the data set is associated with species ID. If one compares the species in Fig. 4B to the phylogeny in Fig. 3 it is immediately clear that the taxa that respond most to selection (and rely on de novo mutations) cluster in the lower part of the tree and are all unicellular organisms.

These results provide food for thought, especially in the light of global changes (Masson-Delmotte *et al*., 2021). The overall positive response to selection that we report indicates that a rather diverse set of species may respond evolutionarily to increased thermal stress. Clearly, unicellular organisms seem to respond more easily by evolution. Of course, we here cannot analyse the speed of the evolutionary response which is a central point when populations have to adapt to potentially rapid environmental change. This is important, because evolutionary changes and adaptation to thermal stress may lead to monopolization and actually prevent interacting species from tracking environmental change, resulting in overall loss of biodiversity, as predicted theoretically by Thompson & Fronhofer (2019).

A cautious interpretation of these selection experiments is also warranted because natural systems are obviously more complex than laboratory populations (but see Buckley & Kingsolver, 2021). Specifically, the studies analysed here mainly use fecundity or population growth rate as a measure of fitness (see Table S1) which can be problematic (for a recent discussion, see Kokko, 2021). Fitness is usually multidimensional with different fitness components such as fecundity and viability reacting differently to thermal selection (see e.g., Wadgymar *et al*., 2017). Under such conditions, where vital rates may trade-off with each other, complex eco-evolutionary feedbacks can be expected under climate change (Cotto *et al*., 2019).

### Caveats of linear reaction norms

Surprisingly, we did not recapture this increase in fitness at the selected temperature when considering studies that had also assay fitness at one additional temperature (Fig. 5). We suggest that this result should not be over-interpreted biologically. Firstly, this part of the meta-analysis has the least power since we only have 61 TPCs from 18 papers with linear reaction norms. Secondly, and most importantly, TPCs are well known to be highly non-linear (discussed in Huey & Kingsolver, 1993, and many others). This suggests that linear reaction norms may not be the most appropriate tool for describing evolutionary responses of a non-linear performance curve. This interpretation gains some weight since we do recapture signals of successful adaptation when analysing data from studies that explore TPC shapes in more detail (Fig. 6).

### Thermal adaptation to higher temperatures trades off with fitness at **lower** temperatures

Analysing responses of better resolved TPCs (here, three and more temperatures assayed; Fig. 6) we recapture similar results compared to the response to selection at the selection temperature (Fig. 4). We observe a clear positive response to selection at the selection temperature and above, as well as the already discussed effect of the type of genetic variation, although the latter is rather weak.

While the exact shape of the function we fit here only serves descriptive purposes, it indicates that, overall, after thermal selection, fitness is increased at and beyond the selection temperature. This comes at a fitness cost immediately below the selection temperature and for extreme temperatures. This asymmetry indicates a potential cost of adaptation to higher temperatures that trades off with fitness at lower temperatures. In the context of Huey & Kingsolver (1993), we therefore find evidence for entire TPC shifts rather than changes in TPC width (see also Fig. 1). This finding is in good accordance with recent work by Logan & Cox (2020) who find that species seem to either adapt to shifting means or to variance but not to both. Here, all studies analysed focus on adaptation to constant temperatures. Analogous results of fitness gains at high temperatures associated with losses at low temperatures have also been observed in ants due to urbanisation-related temperature increases (Diamond *et al*., 2017). The fact that we find adaptation at the selection temperature (Fig. 4) and an increase in performance slightly above (Fig. 6) may hint that TPCs at intermediate temperatures may be more evolvable than suggested by Logan & Cox (2020) who argue that tolerance limits are more readily evolvable based on their review of the literature.

Furthermore, the asymmetry of the response and especially increased fitness beyond the selection temperature (Fig. 6) is in good accordance with Martin & Huey (2008) who argue that due to Jensen’s inequality and the asymmetry of the TPC, the maxima of TPCs should be above the experienced temperature. Finally, the decrease in fitness for low temperatures does not provide support for the “Hotter is wider” hypothesis, at least not in its extreme form as represented in Fig. 1. However, our analysis does not allow us to quantify this effect in detail.

### Limited support for “Hotter is better”

Our data-set also allows us to investigate the “Hotter is Better” hypothesis (Angilletta *et al*., 2010) by comparing fitness at the respective maxima of TPCs before and after selection (Fig. 7). Overall, we did not obtain any strong response supporting “Hotter is Better” although the data indicate a trend in the expected direction. This result should not be over-interpreted as the analysis is rather under-powered with only 27 data points from 11 papers.

### Thermal adaptation in plants

In our systematic search, we only found one study on plants (*Brassica napus*). This lack of studies on plants is in line with recent work by Lancaster & Humphreys (2020) who discuss how limited knowledge is available on thermal niches in plants. Lancaster & Humphreys (2020) provide a major database of thermal tolerance in plants and highlight that macrophysiological rules developed in animals seem to also hold in plants.

Of course, this absence of studies may not be very surprising, especially when considering that some vascular plants and trees have long generation times, and, therefore, genetic adaptation may be expected to be slow. Instead, these long-lived organisms may cope with increasing temperatures via phenotypic plasticity. Plasticity allows for rapid responses to the variable environmental conditions experienced by trees during their long lifespan, and will likely represent their first responses to climate changes (Savolainen *et al*., 2004; Chevin *et al*., 2013; Franks *et al*., 2014). However, such responses may not be rapid enough or adaptive for all species (Duputié *et al*., 2015), and more investigations are clearly needed.

### Thermal adaptation in viruses

In the above presented meta-analysis we chose to exclude viruses because of their unique obligate parasitic strategy which hijacks the cellular DNA or RNA replication machinery of the host (Ryu, 2017). Therefore their efficiency of replication would have to be at least partially dependent on the metabolic effects of temperature on the host enzymes along with its own potential adaptations to a given temperature. Nevertheless, researchers have addressed a range of important questions in evolutionary biology, and thermal adaptation more specifically, using viruses as model systems.

Arribas *et al*. (2014), for example, observed convergent evolution across different temperature selection regimes. In fluctuating temperature selection regimes, these authors report that adaptation occurs to the most stressful temperature in the range of selection temperatures. At the same time, Hao *et al*. (2015) show that fluctuating temperatures may make evolutionary rescue, that is, evolutionary change that leads to the recovery of declining populations (Gonzalez *et al*., 2013), more difficult.

At a more mechanistic level, Betancourt (2009) studied mutations occurring during temperature adaptation and concluded that early emerging mutations had the largest effect on the adaptation to higher temperatures. Poon & Chao (2004) established the increased importance of recombination as populations face genetic drift due to extremely small population sizes when challenged with adaptation to increasing temperature. They used viruses which are able to exchange genetic materials when multiple viruses infect a single host cell.

Temperature adaptation can also have correlated effects in viruses: Carratala *et al*. (2020) show that with adaptation to higher temperatures, viruses also evolved to handle stress induced by free chlorine, making it difficult to eliminate disease-causing viruses in wastewater-treatment plants, for example. With the use of heat shocks, Domingo-Calap *et al*. (2010) demonstrated that populations of viruses which adapted to higher temperatures also showed higher mutational robustness in the presence of mutation inducing chemicals like nitrous acid.

Finally, Knies *et al*. (2009) and Knies *et al*. (2006) addressed the phenomena of “Hotter is better” and “Hotter is wider” by the use of both experimentally and naturally evolved high temperature populations and successfully validated these phenomena in viruses. All of these studies highlight that viruses can also adapt to changing temperatures readily.

### Adaptation to fluctuating temperatures

In our analysis, we have only considered evolutionary responses to constant selection temperatures, but experiments with fluctuating temperature regimes have been performed earlier-on. For example, Bennett *et al*. (1992) have used *Escherichia coli* in an experiment including intergenerational fluctuations between high and low temperatures and showed that the bacteria evolved reduced variability for fitness when assayed across a range of temperatures after selection compared to the thermal specialists evolved at specific constant temperatures. In a further experiment, Lenski & Bennett (1993) showed that even though *E. coli* was selected at fluctuating temperatures, lines adapted asymmetrically more to high than to low temperatures.

In contrast, populations of the ciliate *Paramecium caudatum* facing random daily fluctuations between high and low temperatures exhibited the evolution of a superior generalist which performed better or at least as well as the high and low temperature specialist populations (Duncan *et al*., 2011). Similar results have been reported by Ketola *et al*. (2013) for the bacterium *Serratia marcescens. S. marcescens* temperature generalists also exhibited exaptation to novel environments and stresses at the cost of characteristics like invasiveness and virulence. An increasing amplitude of fluctuation in temperatures can also lead to the evolution of increased temperature optima and maximal growth rates (Bonnefond *et al*., 2017).

However, using the dung fly *Sepsis punctum*, Berger *et al*. (2014) failed to evolve a superior generalist even though they included an intra- and intergenerational temperature fluctuation selection treatment. These authors also did not find any increase in the breadth of the TPC even though these populations showed an overall reduction in maximum fitness compared to the constant temperature selection lines. The results of the intragenerational fluctuation treatment supported the “Hotter is better” hypothesis. In addition, evolution of compensatory mechanisms like heat shock proteins and their expression levels may mask effects of fluctuating temperatures: Ketola *et al*. (2004) showed in the ciliate *Tetrahymena thermophila* that the expression of heat shock protein was highest in rapidly fluctuating environments.

The large variation between the responses to fluctuating temperature selection observed across systems may be due to an erroneous measurement of fitness: Generally, fitness of such populations is assayed at a constant temperatures, however, Ketola & Saarinen (2015) argue that here fitness should be assayed in a fluctuating environmental. Accordingly, Saarinen *et al*. (2018) showed for nine bacterial species that were selected in fluctuating temperatures, that they achieved higher yields when assayed in a fluctuating temperature assay compared to constant conditions.

### Thermal adaptation and biotic interactions

In nature, adaptation to temperature will most likely be taking place in the presence of biotic interactions. Focusing in intra-specific competition, Van Doorslaer *et al*. (2009) exposed *Daphnia magna* populations to varying culling regimes, that is, different levels of intraspecific competition, while applying a thermal selection treatment simultaneously. Populations issued from a harsher culling regime had an overall higher performance at both the low and high assay temperatures when selected at high temperature, compared to populations which experienced milder culling regimes.

Similarly, the presence of a predator can also drastically affect the outcome of thermal selection (Tseng & O’Connor, 2015; Tseng *et al*., 2019). Using *Daphnia pulex*, Tseng & O’Connor (2015) reported that selection at high temperatures only led to increased population growth rates in the presence of a predator. The presence of a predator additionally led to increased TPC width. Effects on thermal plasticity could also be mediated by bottom-up effects linked to the thermal evolution background of resources (Tseng *et al*., 2019).

Biotic interactions may also lead to events of exaptation where adaptation to high temperatures beyond the existing thermal niche can take place without even actively selecting for that particular high temperature. Zhou *et al*. (2019) report that when the yeast *Lachancea thermotolerans* was co-cultured with bacteria, an increase in thermotolerance was observed. Similarly, Alto & Turner (2009) showed that vesicular stomatitis virus adaptation to different hosts exhibited different TPCs: Viruses which were host generalists had a narrower TPC than viruses which were host specialists. The authors attribute this difference to the loss of thermal stability of viral genes and pleiotropic effects between viral genes for thermal adaptation, when adapting to a variety of hosts. In contrast, coevolution between a host and a bacteriophage reduced thermal adaptation in the phage (Zhang & Buckling, 2011).

In summary, biotic interactions affect the evolutionary trajectories of thermal adaptation. Be it intraspecific competition, antagonistic biotic interactions like predation or antagonistic coevolution, they change the impact of temperature stress on the temperature selected organism, either accelerating and delaying adaptation.

## Conclusions

In conclusion, our results clearly show that thermal performance curves (TPCs), across the tree of life, have the potential to evolve in response to temperature selection and not only thermal limits (Logan & Cox, 2020). Our analyses indicate that thermal adaptation leads to overall niche shifts and that adaptation to higher temperature trades off with fitness at lower temperature.

Clearly, these results apply strictly speaking to laboratory systems and, in nature, biotic interactions and fluctuating temperatures may modulate the picture (Buckley & Kingsolver, 2021; Kellermann & van Heerwaarden, 2019). In addition, in nature, interactions between temperature adaptation and adaptation to other abiotic stressors may also play a role (Yilmaz *et al*., 2020). Comprehensively understanding adaptation to multiple stressors is therefore an important challenge, especially given the complexities involved already in a purely ecological context (see e.g. Jackson *et al*., 2021).

We found only one paper which contained a temperature selection experiment with plants which aligns well with the conclusion of Lancaster & Humphreys (2020) that knowledge about TPCs in plants remains very limited. This raises the question whether the general patterns of temperature adaptation discussed above hold in vascular plants other than algae and unicellular photosynthesizing organisms that are represented here. Along similar lines, experiments on larger organisms and ecologically impactful taxa such as marine plankton (Collins *et al*., 2020) remain a minority which critically reduces our ability to infer responses to global warming (see also Sinclair *et al*., 2016). While our work shows that the existing observed responses to thermal selection are encouraging, it is also clear that a larger taxonomic diversity needs to be included in temperature selection experiments in the future.

## Author contributions

E.A.F. and S.P.M. conceived the study. S.P.M. gathered the data. G.Z. performed the statistical analyses. S.P.M. and E.A.F. wrote the manuscript and all authors commented on the draft.

## Acknowledgements

This work was supported by a grant from the Agence Nationale de la Recherche (No.: ANR-19-CE02-0015) to EAF. This is publication ISEM-YYYY-XXX of the Institut des Sciences de l’Evolution – Montpellier.

## Code and Data availability

Data and R-code for the meta-analysis are available from XXX.

