## Supplementary Material for "Evolution of thermal performance curves: a meta-analysis of selection experiments"

### S1 Web of Science search algorithm

ts = (temperature OR thermal)

AND

ts = (experimental evolution OR artificial selection OR selection experiment OR laboratory selection OR adaptation)

AND

ts = (microcosm* OR mesocosm* OR chemostat* OR experiment* OR evolutionary rescue)

NOT

ts=(cancer* OR cardio* OR surg* OR carcin* OR medic* OR drug*)

AND

wc = (Behavioral Sciences OR Ecology OR Evolutionary Biology OR Limnology OR Marine & Freshwater Biology OR Microbiology OR Multidisciplinary Sciences OR Plant Sciences OR Zoology OR Biochemical Research Methods OR Environmental Sciences OR Genetics & Heredity OR Biochemistry & Molecular Biology OR Life Sciences & Biomedicine - Other Topics)

AND

py = 1900-2021

ISI WoS field tag explanation: ts (Topic), wc (Web of Science Categories), py (Year Published)

### S2 Criteria for rejection of papers

1. Ecological papers with transplant experiments, i.e., no selection experiment or experimental evolution performed.

2. Ecological papers measuring the TPC without a selection experiment.

3. Ecological papers in which study organisms have to actively select a thermal environment in a choice experiment, e.g. along a thermal gradient arena, in order to study for example, matching habitat choice.

4. Ecological papers with parental effects of temperature treatment on the offspring and their response which are generally measured over only one generation.

5. Experimental evolution papers with only life-history traits like body size, developmental time, and no direct fitness measure reported.

6. Experimental evolution papers with a direct fitness measure present but the assay temperatures for the selection and the control/ ancestor lines do not correspond making it impossible to calculate relative fitness at a particular assay temperature.

7. Experimental evolution experiments where a control or an ancestor are absent and only the fitness trajectory of the selection line is studied with increasing temperature.

8. Experimental evolution experiments that did not report standard errors or other error estimates.

### S3 Supplementary statistical methods

#### Direct response to selection

We fitted linear mixed models to this data subset comprising all additive models combining the following explanatory variables: relative selection temperature as a categorical explanatory variable (higher, lower or equal compared to the control treatment) and type of genetic variation used (standing genetic variation vs. de novo variation based on mutations). This analysis included 230 data points from 40 studies and 28 species.

We chose vaguely informative priors: the intercept parameter priors for all models followed a normal distribution with mean 0 and standard deviation 2. The mean and standard deviation of the random effect hyperpriors followed a normal distribution with mean 0 and standard deviation 3.

We further ran a supplementary analysis using the same basic model set but adding the number of generations an experiment lasted as another explanatory variable. For this analysis, we standardised the number of generations to avoid divergent transitions and removed four studies (seven data points) that did not report the number of generations (normal prior mean 0 and standard deviation 1). The additional analysis included 223 data points from 36 studies and 26 species.

#### Two-point analysis

In analogy to the analysis of the direct response to selection, we fitted linear mixed models explaining relative fitness as a function of (absolute) relative assay temperature. In the full model, we allowed for different slopes and intercepts depending on whether experimental populations had evolved at higher or lower temperature compared to the control and tested in the final assay at high or low temperature. We subsequently included an additional explanatory variable, the type of genetic variation, as varying intercept, slope, both intercept and slope. The intercept and slope parameter priors for all models followed a normal distribution with mean 0 and standard deviation 1. The mean and standard deviation of the random effect hyperpriors followed a normal distribution with mean 0 and standard deviation 1. This analysis included 122 data points in 61 TPCs from 18 studies and 15 species.

A supplementary analysis was conducted using the same model details but adding the standardized number of generations as additional explanatory variable. Due to missing data on the number of generations, we removed 10 TPCs from this additional analysis. This analysis included 102 data points in 51 TPCs from 13 studies and 12 species.

#### Multipoint analysis

Here, we first fitted mixed models including intercept-only, linear, quadratic and cubic models of relative fitness as a function of standardized relative assay temperature. Note that these shapes are descriptive and flexible enough to describe the shapes we find. We do not imply that TPCs are actually cubic. All priors followed a normal distribution with mean 0 and standard deviation 1. The mean and standard deviation of the random effect hyperpriors followed a normal distribution with mean 0 and standard deviation 1. This analysis included 600 data points in 159 TPCs from 21 studies and 13 species.

We also added the type of genetic variation as an explanatory variable. We fitted supplementary models with the inclusion of the number of generations (standardized) as an extra explanatory variable. Due to missing data on the number of generations, we removed 3 TPCs from this additional analysis. This analysis included 586 data points in 156 TPCs from 19 studies and 11 species.

#### "Hotter is better"

Similar to the direct response to selection analysis (see above), we fitted two linear mixed models: an intercept model to test whether the difference in maximal relative fitness was different from zero and a second one in which the temperature selection regime (selection at higher or lower temperatures than the ancestor/ control) was added as a categorical fixed effect. In both models, the priors for the intercepts followed a normal distribution with mean 0 and standard deviation 2. The random effect hyperprior mean and standard deviation followed a normal distribution with mean 0 and standard deviation 3. This analysis included 27 data points from 11 studies and 9 species.

We then added to this model the extra explanatory variable type of genetic variation as varying intercept. We conducted a supplementary analysis further considering number of generations (standardized) as varying slope. Here, we removed one study not reporting this information. This analysis included 26 data points from 10 studies and 8 species.

#### Assessment of publication bias

We assessed publication bias using funnel plots (Sterne *et al.*, 2001; Sterne & Egger, 2001; O'Dea *et al.*, 2021), where the inverse of the standard error is plotted against relative fitness. In the absence of bias, funnel plots are expected to be symmetrical. In addition to visual inspection, symmetry can be inspected by means of linear regression of the two variables. The regression line is expected to intercept the x-axis close to the origin if there is bias. Alternatively, if the studies are evenly distributed and thus there is no bias towards high or low relative fitness values, a zero-slope (horizontally flat line) may be expected.

We inspected the symmetry not only visually, but we also tested and quantified it by fitting an intercept model and a linear regression model (chain length: warmup = 1,000 iterations, chain = 30,000 iterations). The intercept and slope priors for both models followed a normal distribution with mean 0 and standard deviation 1. Relative fitness was standardised in cases of divergent transitions. We included species ID, study ID and TPC ID as random effects as in the main analyses. The mean and standard deviation of the random effect hyperpriors followed a normal distribution with mean 0 and standard deviation 1. Based on WAIC weights we averaged the posterior predictions of the models.

#### S4 Supplementary figures

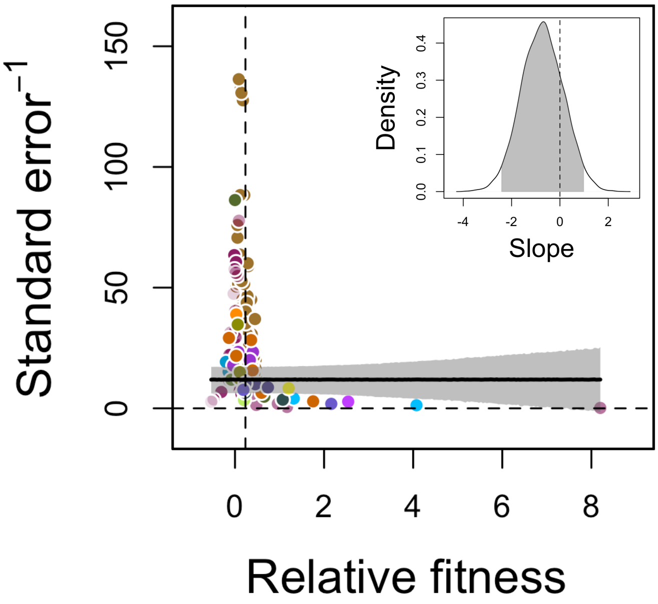


Figure S1: Funnel plot of the analysis of assay data measured at the same temperature experienced during experimental evolution. Each point is a study with different colours per species (color code see Fig. 3). The shaded area and thick line are the overall 95% compatibility interval and median of the averaged posterior predictions based on WAIC weights. Model selection results are reported in Table S3. The random effect structure used for the corresponding main analysis was included here. The insert panel highlights the posterior distribution of the slope parameter which clearly overlaps zero.

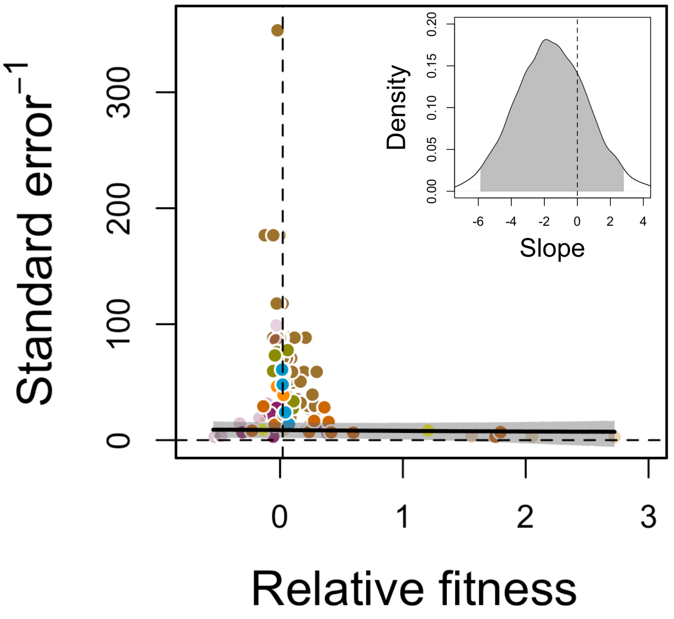


Figure S2: Funnel plot of the two temperature assay point analysis. Each point is a study with different colours per species (color code see Fig. 3). The shaded area and thick line are the overall 95% compatibility interval and median of the averaged posterior predictions based on WAIC weights. Model selection results are reported in Table S4. The random effect structure used for the corresponding main analysis was included here. The insert panel highlights the posterior distribution of the slope parameter which clearly overlaps zero.

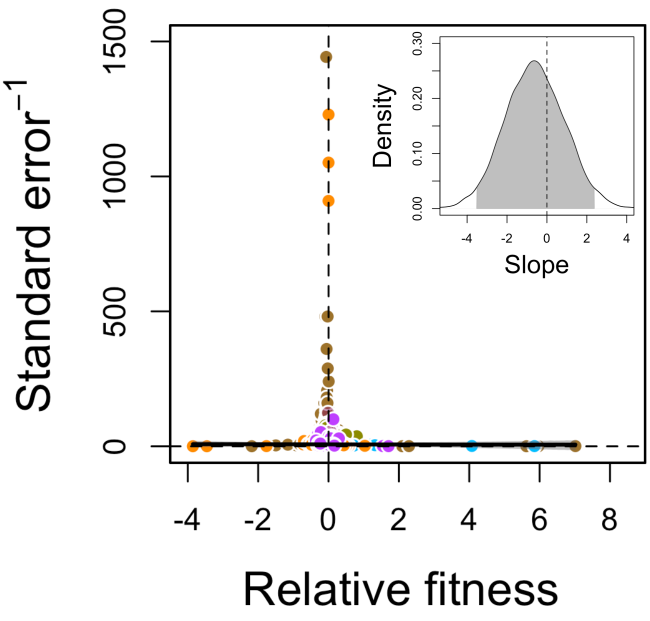


Figure S3: Funnel plot of the multipoint temperature assay analysis. Each point is a study with different colours per species (color code see Fig. 3). The shaded area and thick line are the overall 95% compatibility interval and median of the averaged posterior predictions based on WAIC weights. Model selection results are reported in Table S5. The random effect structure used for the corresponding main analysis was included here. The insert panel highlights the posterior distribution of the slope parameter which clearly overlaps zero.

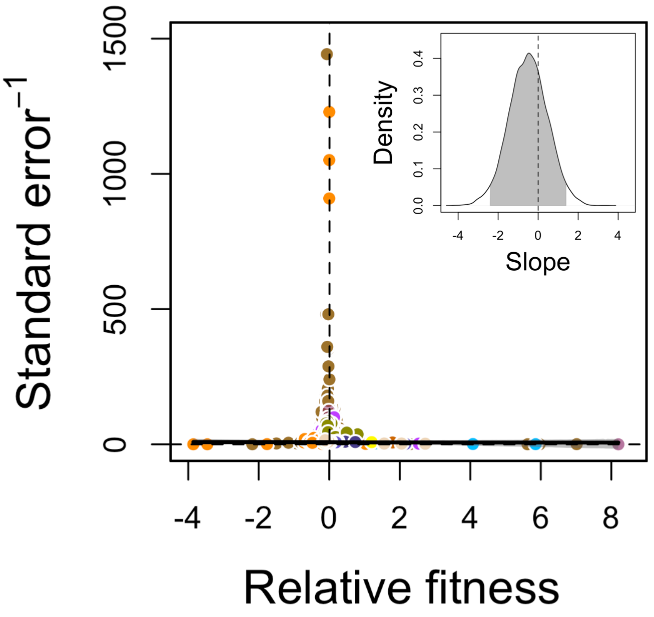


Figure S4: Funnel plot of all data together, including one, two, and multipoint temperature assay analysis. Each point is a study with different colours per species (color code see Fig. 3). The shaded area and thick line are the overall 95% compatibility interval and median of the averaged posterior predictions based on WAIC weights. Model selection results are reported in Table S6. The random effect structure used for the corresponding main analysis was included here. The insert panel highlights the posterior distribution of the slope parameter which clearly overlaps zero.

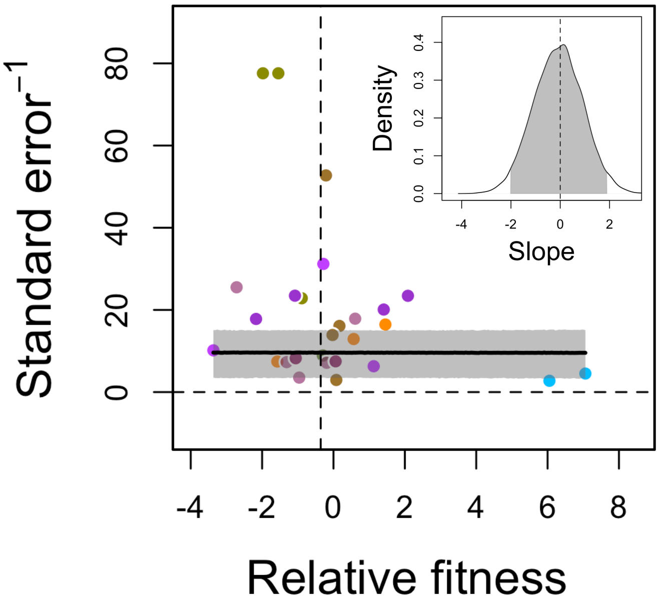


Figure S5: Funnel plot of the "Hotter is better" analysis. Each point is a study with different colours per species (color code see Fig. 3). The shaded area and thick line are the overall 95% compatibility interval and median of the averaged posterior predictions based on WAIC weights. Model selection results are reported in Table S7. The random effect structure used for the corresponding main analysis was included here. The insert panel highlights the posterior distribution of the slope parameter which clearly overlaps zero.

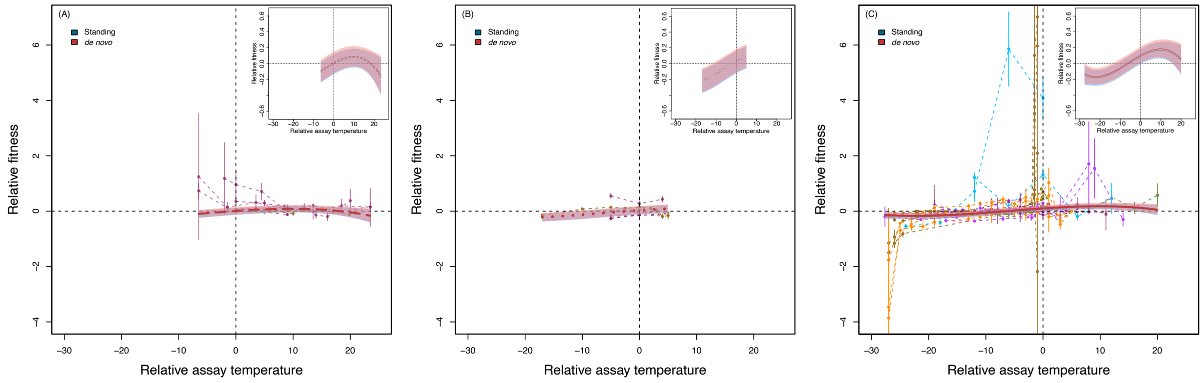


Figure S6: Evolution of the TPC. The plot shows relative fitness as a function of relative assay temperature, that is, the difference between selection and assay temperature, for studies that assayed selected lines at three and more temperatures allowing us to infer changes in TPC shape. Each combination of points connected by a dashed line corresponds to one TPC and the error bars represent the associated standard error. Different colours correspond to different species as shown in Fig. 3. The solid lines visualize the median model predictions and the shaded areas are the 95% compatibility interval. Here, the best model (Tables S12 and S13) is a cubic model model that includes an effect of the type of genetic variation used in the experiments (standing genetic variation vs. de novo mutations; represented in blue and red, respectively) and relative selection temperature ('Sign'), that is, whether selection happened at lower (A), equal (B) or higher (C) temperatures compared to the control treatment. The insets show only the statistical model prediction for better visibility of the compatibility intervals.

#### S5 Supplementary tables

Table S1: Overview of selected papers, with taxonomic information, fitness measure used, source of data, as well as details on the calculation of relative fitness which is described in the main text. Equation numbers refer to Chevin (2011).

| **Reference** | **Species** | **Fitness measure** | **Data source** | **Relative fitness calculation** |
|---|---|---|---|---|
| Cullum *et al.* (2001) | *Escherichia coli* | Relative fitness | Paper | Eq. 3.2 |
| Dam *et al.* (2021) | *Acartia tonsa* | Population fitness | Graph | Eq. 3.2 |
| Barton *et al.* (2020) | *Ostreococcus tauri* | Growth rate | SI | Eq. 3.2 |
| Frenck *et al.* (2013) | *Brassica napus* | Seed quantity | Graph | Eq. 2.5 |
| Gomez-Llano *et al.* (2021) | *Drosophila melanogaster* | Number of offspring produced | Graph | Eq. 2.5 |
| Rouco *et al.* (2011) | *Microcystis aeruginosa* | Doubling time | Graph | Eq. 3.2 |
| Sandberg *et al.* (2014) | *Escherichia coli* | Population growth rate | Paper | Eq. 3.2 |
| Wu *et al.* (2020) | *Phytophthora infestans* | Colony size | SI | Eq. 2.5 |
| Bennett & Lenski (2007) | *Escherichia coli* | Relative fitness | Paper | Eq. 3.2 |
| Zhang *et al.* (2019) | *Daphnia magna* | Intrinsic rate of increase | SI | Eq. 2.5 |
| Chakravarti *et al.* (2017) | *Symbiodinium* sp. | Specific growth rate | SI | Eq. 3.2 |
| Duncan *et al.* (2011) | *Paramecium caudatum* | Area under the growth curve | Author | Eq. 3.2 |
| Van Doorslaer *et al.* (2009) | *Daphnia magna* | Performance | Graph | Eq. 2.5 |
| Plesnar-Bielak *et al.* (2012) | *Rhizoglyphus robini* | Fecundity | Graph | Eq. 2.5 |
| Partridge *et al.* (1995) | *Drosophila melanogaster* | Fecundity | Paper | Eq. 2.5 |
| Tseng & O'Connor (2015) | *Daphnia pulex* | Per capita growth rate | Graph | Eq. 2.5 |
| Hallsson & Björklund (2012) | *Callosobruchus maculatus* | Fecundity | Graph | Eq. 2.5 |
| Shi & Xia (2003) | *Pseudomonas pseudoalcaligenes* | Maximum growth rate | Paper | Eq. 3.2 |
| Batarseh *et al.* (2020) | *Escherichia coli* | Relative fitness | SI | Eq. 3.2 |
| Koch & Guillaume (2020) | *Tribolium castaneum* | Offspring number | Graph | Eq. 2.5 |
| Lind *et al.* (2020) | *Caenorhabditis remanei* | Daily growth factor | Graph | Eq. 2.5 |
| Schluter *et al.* (2014) | *Emiliania huxleyi* | Exponential growth rate | Graph | Eq. 3.2 |
| Tarkington & Zufall (2021) | *Tetrahymena thermophila* | Relative fitness | Paper | Eq. 3.2 |
| Van Doorslaer *et al.* (2010) | *Daphnia magna* | Growth rate | Graph | Eq. 2.5 |
| Breckels *et al.* (2014) | *Poecilia reticulata* | Brood size | Paper | Eq. 2.5 |
| Xu (2004) | *Cryptococcus neoformans* | Relative fitness colony size | Paper | Eq. 2.5 |
| Bennett *et al.* (1992) | *Escherichia coli* | Relative fitness | Paper data | Eq. 3.2 |
| Santos (2007) | *Drosophila subobscura* | Average relative fitness | Graph | Eq. 2.5 |
| Van Doorslaer *et al.* (2007) | *Simocephalus vetulus* | Performance | Graph | Eq. 3.2 |
| Berger *et al.* (2014) | *Sepsis punctum* | Growth rate | SI | Eq. 2.5 |
| Rodriguez-Verdugo *et al.* (2014) | *Escherichia coli* | Relative fitness | SI | Eq. 3.2 |
| Fragata *et al.* (2016) | *Drosophila subobscura* | Fecundity | SI | Eq. 2.5 |
| Pfenninger & Foucault (2020) | *Chironomus riparius* | Population growth rate | Author | Eq. 2.5 |
| Santos *et al.* (2006) | *Drosophila subobscura* | Viability | Graph | Eq. 2.5 |
| Schaum *et al.* (2017) | *Chlamydomonas reinhardtii* | ln mu_max | Graph | Eq. 3.2 |
| Cooper *et al.* (2001) | *Escherichia coli* | Maximum growth rate | Graph | Eq. 3.2 |
| Mongold *et al.* (1996) | *Escherichia coli* | Relative fitness | Graph | Eq. 3.2 |
| Killeen *et al.* (2017) | *Paramecium caudatum* | Growth rate | SI | Eq. 3.2 |
| O'Donnell *et al.* (2018) | *Thalassiosira pseudonana* | Specific growth rate | SI | Eq. 3.2 |
| Gilchrist *et al.* (1997) | *Drosophila melanogaster* | Walking speed | Graph | Eq. 2.5 |
| Listmann *et al.* (2016) | *Emiliania huxleyi* | Exponential growth rate | SI | Eq. 3.1 |
| Tseng *et al.* (2019) | *Daphnia pulex* | Per capita growth rate | SI | Eq. 2.5 |
| Aranguren-Gassis *et al.* (2019) | *Chaetoceros simplex* | Growth rate | SI | Eq. 3.1 |
| Condon *et al.* (2014) | *Drosophila melanogaster* | Fecundity | Graph | Eq. 2.5 |
| Romero-Olivares *et al.* (2015) | *Neurospora discreta* | Mycelial growth rate | Graph | Eq. 2.5 |
| Schaum *et al.* (2018) | *Thalassiosira pseudonana* | ln(growth rate) | Graph | Eq. 3.2 |
| Mongold *et al.* (1999) | *Escherichia coli* | Absolute fitness | Graph | Eq. 3.1 |

Table S2: Number of studies and the number of TPCs associated with each organism for all selected studies of the meta-analysis.

| **Species** | **No. of studies** | **No. of TPCs** |
|---|---|---|
| *Daphnia magna* | 3 | 5 |
| *Daphnia pulex* | 3 | 4 |
| *Simocephalus vetulus* | 1 | 2 |
| *Acartia tonsa* | 1 | 3 |
| *Drosophila subobscura* | 3 | 8 |
| *Drosophila melanogaster* | 4 | 9 |
| *Sepsis punctum* | 1 | 1 |
| *Chironomus riparius* | 1 | 2 |
| *Tribolium castaneum* | 1 | 1 |
| *Callosobruchus maculatus* | 1 | 1 |
| *Rhizoglyphus robini* | 1 | 2 |
| *Caenorhabditis remanei* | 1 | 2 |
| *Poecilia reticulata* | 1 | 1 |
| *Cryptococcus neoformans* | 1 | 2 |
| *Neurospora discreta* | 1 | 2 |
| *Ostreococcus tauri* | 1 | 2 |
| *Chlamydomonas reinhardtii* | 1 | 1 |
| *Brassica napus* | 1 | 1 |
| *Chaetoceros simplex* | 1 | 4 |
| *Thalassiosira pseudonana* | 3 | 5 |
| *Phytophthora infestans* | 1 | 2 |
| *Tetrahymena thermophila* | 1 | 6 |
| *Paramecium caudatum* | 2 | 4 |
| *Symbiodinium* sp. | 1 | 1 |
| *Emiliania huxleyi* | 2 | 6 |
| *Microcystis aeruginosa* | 1 | 3 |
| *Synechococcus* sp. | 1 | 2 |
| *Pseudomonas pseudoalcaligenes* | 1 | 3 |
| *Escherichia coli* | 9 | 164 |

Table S3: Model selection table for analysis of the overall effect of selection funnel plot (fitness data measured at the same temperature as experienced during selection). We report WAIC, the standard error of WAIC ('SE'), the WAIC different to the bets model (dWAIC) as well as the WAIC weights.

| **Model** | **WAIC** | **SE** | **dWAIC** | **weight** |
|---|---|---|---|---|
| intercept | 1940.3 | 37.52 | 0 | 0.59 |
| intercept + slope | 1941.0 | 37.52 | 0.7 | 0.41 |

Table S4: Model selection table for two-point analysis funnel plot. We report WAIC, the standard error of WAIC ('SE'), the WAIC different to the bets model (dWAIC) as well as the WAIC weights.

| **Model** | **WAIC** | **SE** | **dWAIC** | **weight** |
|---|---|---|---|---|
| intercept + slope | 1287.9 | 65.56 | 0.0 | 0.56 |
| intercept | 1288.4 | 65.37 | 0.5 | 0.44 |

Table S5: Model selection table for multi-point analysis funnel plot. We report WAIC, the standard error of WAIC ('SE'), the WAIC different to the bets model (dWAIC) as well as the WAIC weights.

| **Model** | **WAIC** | **SE** | **dWAIC** | **weight** |
|---|---|---|---|---|
| intercept + slope | 7534.4 | 284.39 | 0.0 | 0.53 |
| intercept | 7534.6 | 284.38 | 0.2 | 0.47 |

Table S6: Model selection table for the funnel plot representing all data pooled. We report WAIC, the standard error of WAIC ('SE'), the WAIC different to the bets model (dWAIC) as well as the WAIC weights.

| **Model** | **WAIC** | **SE** | **dWAIC** | **weight** |
|---|---|---|---|---|
| intercept | 9278.1 | 328.62 | 0.0 | 0.52 |
| intercept + slope | 9278.3 | 329.05 | 0.2 | 0.48 |

Table S7: Model selection table for the "Hotter is better" analysis. We report WAIC, the standard error of WAIC ('SE'), the WAIC different to the bets model (dWAIC) as well as the WAIC weights.

| **Model** | **WAIC** | **SE** | **dWAIC** | **weight** |
|---|---|---|---|---|
| intercept | 250.4 | 23.92 | 0.0 | 0.51 |
| intercept + slope | 250.4 | 23.96 | 0.1 | 0.49 |

Table S8: Model selection table for the overall response to thermal selection. Explanatory variables are ‘Type of genetic variation’ (standing genetic variation vs. de novo variation based on mutations; ‘Var’) and ‘Relative selection temperature’ (higher, lower or equal compared to the control treatment; ‘Sign’). In the table ‘+’ indicates an additive effect. We report WAIC, the standard error of WAIC (‘SE’), the WAIC different to the bets model (dWAIC) as well as the WAIC weights.

| Model | WAIC | SE | dWAIC | weight |
|---|---|---|---|---|
| Var | 1623663799 | 485755802 | 0 | 1 |
| Var + Sign | 1714151640 | 513056393 | 90487841 | 0 |
| Intercept | 1782783610 | 534129409 | 159119811 | 0 |
| Sign | 1930577420 | 578697354 | 306913621 | 0 |

Table S9: Model selection table for the overall response to thermal selection using the reduced data set to included number of generations in the analysis. Explanatory variables are ‘Type of genetic variation’ (standing genetic variation vs. de novo variation based on mutations; ‘Var’), ‘Relative selection temperature’ (higher, lower or equal compared to the control treatment; ‘Sign’) and number of generations in the experiment (‘Gen’; standardized). In the table ‘+’ indicates an additive effect and ‘:’ an interactive effect. We report WAIC, the standard error of WAIC (‘SE’), the WAIC different to the bets model (dWAIC) as well as the WAIC weights.

| Model | WAIC | SE | dWAIC | weight |
|---|---|---|---|---|
| Var | 1711459014 | 523308177 | 0 | 1 |
| Gen + Var:Gen | 1737972889 | 531378488 | 26513874 | 0 |
| Var + Gen | 1741041679 | 532422388 | 29582665 | 0 |
| Intercept | 1776068393 | 543640762 | 64609379 | 0 |
| Var + Gen + Var:Gen | 1792392991 | 548092410 | 80933976 | 0 |
| Var + Sign | 1819269616 | 556466483 | 107810601 | 0 |
| Var + Gen + Var:Gen + Sign | 1844078940 | 564070080 | 132619926 | 0 |
| Gen | 1858258633 | 568885469 | 146799619 | 0 |
| Gen + Var:Gen + Sign | 1861953281 | 569617448 | 150494267 | 0 |
| Sign | 1886799538 | 577675125 | 175340524 | 0 |
| Var + Gen + Sign | 1895395580 | 579802412 | 183936565 | 0 |
| Sign + Gen | 1925653718 | 589701442 | 214194704 | 0 |

Table S10: Model selection table for the two-point response to thermal selection analysis. Explanatory variables are absolute relative assay temperature (abbreviated as 'Temp' in the table), sign of the assay temperature, that is, whether the second assay was carried out above, at, or below the selection temperature (abbreviated as 'Sign' in the table) and type of genetic variation ('Var'). Note that in the analysed studies 'Sign' correlates with the temperature of the assay: studies that performed selection at the same or lower temperatures in comparison to the control always performed the second assay at higher temperatures compared to the first assay point, for studies selecting at higher temperatures than the control the assay was always done at temperatures below the first assay point. In the table '+' indicates an additive effect and ':' an interactive effect. We report WAIC, the standard error of WAIC ('SE'), the WAIC different to the bets model (dWAIC) as well as the WAIC weights.

| **Model** | **WAIC** | **SE** | **dWAIC** | **weight** |
|---|---|---|---|---|
| Intercept | 23614381787 | 15324934662 | 0 | 1 |
| Temp + Var:Temp | 23717825649 | 15392349383 | 103443862 | 0 |
| Temp | 23757133779 | 15417003258 | 142751992 | 0 |
| Var | 24145184487 | 15671246974 | 530802700 | 0 |
| Temp + Var | 24194571950 | 15702742522 | 580190163 | 0 |
| Sign | 24558076629 | 15939269068 | 943694842 | 0 |
| Var + Sign | 24590169935 | 15961464453 | 975788148 | 0 |
| Temp + Sign | 24724264320 | 16046580988 | 1109882532 | 0 |
| Temp + Var + Var:Temp | 24884526473 | 16152985021 | 1270144686 | 0 |
| Temp + Sign + Var:Temp | 25003952581 | 16229373382 | 1389570794 | 0 |
| Temp + Var + Var:Temp + Sign | 25275487420 | 16406777226 | 1661105632 | 0 |
| Temp + Var + Sign | 25543644950 | 16580953259 | 1929263162 | 0 |

Table S11: Model selection table for the two-point response to thermal selection analysis with additional explanatory variables including number of generations (reduced data set). Explanatory variables are absolute relative assay temperature (abbreviated as 'Temp' in the table), Type of genetic variation ('Var'), sign of the assay temperature, that is, whether the second assay was carried out above, at, or below the selection temperature (abbreviated as 'Sign' in the table) and the number of generations the experiment lasted ('Gen'; standardized). In the table '+' indicates an additive effect and ':' an interactive effect. We report WAIC, the standard error of WAIC ('SE'), the WAIC different to the bets model (dWAIC) as well as the WAIC weights.

| **Model** | **WAIC** | **SE** | **dWAIC** | **weight** |
|---|---|---|---|---|
| Intercept | 6354484583 | 4121821808 | 0 | 1 |
| Gen | 6391495482 | 4144758709 | 37010899 | 0 |
| Gen + Temp | 6394900591 | 4146133403 | 40416008 | 0 |
| Temp | 6401329257 | 4151651200 | 46844674 | 0 |
| Temp + Temp:Var | 6445826040 | 4180753202 | 91341457 | 0 |
| Sign | 6466283750 | 4194914902 | 111799167 | 0 |
| Gen + Temp + Temp:Var | 6470951669 | 4196199910 | 116467086 | 0 |
| Var | 6518247622 | 4230464961 | 163763039 | 0 |
| Temp + Var | 6590587951 | 4276767932 | 236103368 | 0 |
| Temp + Var + Gen | 6605700412 | 4285507555 | 251215829 | 0 |
| Temp + Temp:Var + Sign | 6610511656 | 4288613918 | 256027073 | 0 |
| Temp + Sign | 6616766453 | 4292732031 | 262281870 | 0 |
| Var + Gen | 6654040956 | 4318228947 | 299556373 | 0 |
| Var + Sign | 6663301132 | 4325674689 | 308816549 | 0 |
| Var + Temp + Temp:Var | 6687473314 | 4340281054 | 332988731 | 0 |
| Var + Temp + Temp:Var + Gen | 6691804412 | 4341784043 | 337319829 | 0 |
| Temp + Var + Sign | 6770683578 | 4394652256 | 416198995 | 0 |
| Temp + Var + Temp:Var + Sign | 6826309994 | 4430331707 | 471825411 | 0 |

Table S12: Model selection table for the multi-point response to thermal selection analysis. Explanatory variables are absolute relative assay temperature (abbreviated as 'Temp' in the table), Type of genetic variation ('Var') and 'Relative selection temperature' (higher, lower or equal compared to the control treatment; 'Sign'). In the table '+' indicates an additive effect and ':' an interactive effect. We report WAIC, the standard error of WAIC ('SE'), the WAIC different to the bets model (dWAIC) as well as the WAIC weights.

| **Model** | **WAIC** | **SE** | **dWAIC** | **weight** |
|---|---|---|---|---|
| Temp + $Temp^2$ + $Temp^3$ + Var + Sign | 5.202676e+12 | 2.676116e+12 | 0 | 1 |
| Temp + $Temp^2$ + $Temp^3$ + Sign | 5.224529e+12 | 2.687450e+12 | 2.185324e+10 | 0 |
| Temp + $Temp^2$ + $Temp^3$ | 5.290472e+12 | 2.721611e+12 | 8.779641e+10 | 0 |
| Temp + $Temp^2$ + $Temp^3$ + Var | 5.305165e+12 | 2.728286e+12 | 1.024890e+11 | 0 |
| Temp + $Temp^2$ + Sign | 5.621206e+12 | 2.893259e+12 | 4.185308e+11 | 0 |
| Temp + Sign | 5.653209e+12 | 2.909017e+12 | 4.505335e+11 | 0 |
| Temp + $Temp^2$ + Var | 5.683915e+12 | 2.926735e+12 | 4.812390e+11 | 0 |
| Temp + Var + Sign | 5.706087e+12 | 2.937974e+12 | 5.034112e+11 | 0 |
| Temp + $Temp^2$ + Var + Sign | 5.710847e+12 | 2.939725e+12 | 5.081711e+11 | 0 |
| Temp + $Temp^2$ | 5.745498e+12 | 2.956929e+12 | 5.428220e+11 | 0 |
| Temp | 5.821055e+12 | 2.995173e+12 | 6.183796e+11 | 0 |
| Temp + Var | 5.888742e+12 | 3.031702e+12 | 6.860665e+11 | 0 |
| Intercept | 7.447238e+12 | 3.806424e+12 | 2.244562e+12 | 0 |
| Var + Sign | 7.528571e+12 | 3.847949e+12 | 2.325895e+12 | 0 |
| Sign | 7.543139e+12 | 3.856617e+12 | 2.340464e+12 | 0 |
| Var | 7.553221e+12 | 3.861681e+12 | 2.350545e+12 | 0 |

Table S13: Model selection table for the multi-point response to thermal selection analysis with additional explanatory variables including the number of generations (reduced data set). Explanatory variables are absolute relative assay temperature (abbreviated as 'Temp' in the table), Type of genetic variation ('Var'), 'Relative selection temperature' (higher, lower or equal compared to the control treatment; 'Sign') and the number of generations the experiment lasted ('Gen'; standardized). In the table '+' indicates an additive effect and ':' an interactive effect. We report WAIC, the standard error of WAIC ('SE'), the WAIC different to the bets model (dWAIC) as well as the WAIC weights.

| **Model** | **WAIC** | **SE** | **dWAIC** | **weight** |
|---|---|---|---|---|
| Temp + $Temp^2$ + $Temp^3$ + Var + Sign | 5.088572e+12 | 2.615790e+12 | 0 | 1 |
| Temp + $Temp^2$ + $Temp^3$ + Var + Sign + Gen | 5.115694e+12 | 2.627099e+12 | 2.712214e+10 | 0 |
| Temp + $Temp^2$ + $Temp^3$ | 5.127109e+12 | 2.633056e+12 | 3.853778e+10 | 0 |
| Temp + $Temp^2$ + $Temp^3$ + Sign | 5.132144e+12 | 2.637492e+12 | 4.357262e+10 | 0 |
| Temp + $Temp^2$ + $Temp^3$ + Sign + Gen | 5.170180e+12 | 2.656259e+12 | 8.160809e+10 | 0 |
| Temp + $Temp^2$ + $Temp^3$ + Var + Gen | 5.208339e+12 | 2.676779e+12 | 1.197676e+11 | 0 |
| Temp + $Temp^2$ + $Temp^3$ + Gen | 5.218445e+12 | 2.680827e+12 | 1.298730e+11 | 0 |
| Temp + $Temp^2$ + $Temp^3$ + Var | 5.229531e+12 | 2.687119e+12 | 1.409589e+11 | 0 |
| Temp + Sign | 5.466533e+12 | 2.810131e+12 | 3.779613e+11 | 0 |
| Temp + $Temp^2$ + Sign | 5.487874e+12 | 2.821809e+12 | 3.993027e+11 | 0 |
| Temp + $Temp^2$ + Var + Sign | 5.506660e+12 | 2.831523e+12 | 4.180887e+11 | 0 |
| Temp + $Temp^2$ + Gen | 5.513372e+12 | 2.834466e+12 | 4.248003e+11 | 0 |
| Temp + Var + Sign | 5.519486e+12 | 2.838140e+12 | 4.309148e+11 | 0 |
| Temp + Sign + Gen | 5.524629e+12 | 2.841323e+12 | 4.360570e+11 | 0 |
| Temp + Gen | 5.544125e+12 | 2.850432e+12 | 4.555530e+11 | 0 |
| Temp + $Temp^2$ + Sign + Gen | 5.545664e+12 | 2.850498e+12 | 4.570923e+11 | 0 |
| Temp | 5.566529e+12 | 2.862299e+12 | 4.779578e+11 | 0 |
| Temp + $Temp^2$ + Var | 5.583444e+12 | 2.869222e+12 | 4.948724e+11 | 0 |
| Temp + Var + Sign + Gen | 5.595266e+12 | 2.875681e+12 | 5.066942e+11 | 0 |
| Temp + Var + Gen | 5.596104e+12 | 2.877852e+12 | 5.075325e+11 | 0 |
| Temp + Var | 5.611074e+12 | 2.886170e+12 | 5.225020e+11 | 0 |
| Temp + $Temp^2$ + Var + Sign + Gen | 5.617917e+12 | 2.886741e+12 | 5.293456e+11 | 0 |
| Temp + $Temp^2$ | 5.621897e+12 | 2.888791e+12 | 5.333249e+11 | 0 |
| Temp + $Temp^2$ + Var + Gen | 5.623710e+12 | 2.890134e+12 | 5.351387e+11 | 0 |
| Var | 7.316852e+12 | 3.736721e+12 | 2.228280e+12 | 0 |
| Sign | 7.349525e+12 | 3.752395e+12 | 2.260953e+12 | 0 |
| Intercept | 7.365263e+12 | 3.758234e+12 | 2.276691e+12 | 0 |
| Gen | 7.393232e+12 | 3.774242e+12 | 2.304660e+12 | 0 |
| Var + Sign | 7.438562e+12 | 3.797575e+12 | 2.349991e+12 | 0 |
| Sign + Gen | 7.445837e+12 | 3.801358e+12 | 2.357266e+12 | 0 |
| Var + Gen | 7.488237e+12 | 3.823640e+12 | 2.399665e+12 | 0 |
| Var + Sign + Gen | 7.598070e+12 | 3.880620e+12 | 2.509498e+12 | 0 |

Table S14: Model selection table for the "hotter is better" analysis. Explanatory variables are 'Type of genetic variation' (standing genetic variation vs. de novo variation based on mutations) and 'Relative selection temperature' (higher, lower or equal compared to the control treatment). In the table '+' indicates an additive effect. We report WAIC, the standard error of WAIC ('SE'), the WAIC different to the bets model (dWAIC) as well as the WAIC weights.

| **Model** | **WAIC** | **SE** | **dWAIC** | **weight** |
|---|---|---|---|---|
| Intercept | 8660144 | 4963707 | 0 | 1 |
| Sign | 9131870 | 5239167 | 471726.2 | 0 |
| Var | 10781844 | 6203460 | 2121700.4 | 0 |
| Var + Sign | 10785915 | 6205959 | 2125770.9 | 0 |

Table S15: Model selection table for the "hotter is better" analysis using the reduced data set to included number of generations in the analysis. Explanatory variables are 'Type of genetic variation' (standing genetic variation vs. de novo variation based on mutations), 'Relative selection temperature' (higher, lower or equal compared to the control treatment) and the number of generations the experiment lasted ('Gen'; standardized). In the table '+' indicates an additive effect and ':' an interactive effect. We report WAIC, the standard error of WAIC ('SE'), the WAIC different to the bets model (dWAIC) as well as the WAIC weights.

| **Model** | **WAIC** | **SE** | **dWAIC** | **weight** |
|---|---|---|---|---|
| Intercept | 9814054 | 5627564 | 0 | 1 |
| Sign | 10399067 | 5969854 | 585012.6 | 0 |
| Var | 12218321 | 7032109 | 2404267.5 | 0 |
| Var + Sign | 12414274 | 7146141 | 2600220.1 | 0 |
| Gen | 14198806 | 8196894 | 4384751.6 | 0 |
| Gen + Sign | 14539062 | 8395046 | 4725008.6 | 0 |
| Gen + Gen:Var | 15499399 | 8957579 | 5685344.6 | 0 |
| Gen + Gen:Var + Sign | 15659298 | 9053435 | 5845243.5 | 0 |
| Var + Gen | 15703762 | 9078577 | 5889707.6 | 0 |
| Var + Gen + Sign | 15727338 | 9093349 | 5913283.9 | 0 |
| Var + Gen + Var:Gen | 16018136 | 9262200 | 6204082.3 | 0 |
| Var + Gen + Var:Gen + Sign | 16169460 | 9349567 | 6355405.5 | 0 |
